# Pericyte dysfunction and impaired vasomotion are hallmarks of islets during the pathogenesis of type 1 diabetes

**DOI:** 10.1101/2023.03.02.530808

**Authors:** Luciana Mateus Gonçalves, Mirza Muhammad Fahd Qadir, Maria Boulina, Madina Makhmutova, Joana Almaça

**Author notes:** Correspondence to Joana Almaça.

## Abstract

Pancreatic islets are endocrine organs that depend on their microvasculature to function properly. Along with endothelial cells, pericytes comprise the islet microvascular network. These mural cells are crucial for microvascular stability and function, but it is not known if/how they are affected during the development of type 1 diabetes (T1D). Here we investigated islet pericyte density, phenotype and function using living pancreas slices from donors without diabetes, donors with a single T1D-associated autoantibody (Aab+; all GADA+) and recent onset T1D cases. Our data show that islet pericyte and capillary responses to vasoactive stimuli are impaired early on in T1D. Microvascular dysfunction is associated with a switch in the phenotype of islet pericytes towards pro-fibrotic myofibroblasts. Using publicly available RNAseq data, we further found that transcriptional alterations related to endothelin-1 signaling, vascular and ECM remodeling are hallmarks of single Aab+ donor pancreata. Our data show that islet pericyte/microvascular dysfunction is present at early stages of islet autoimmunity.

**Highlights:**

- Changes in islet pericyte coverage and phenotype occur during T1D progression.
- Vascular responses to vasoactive stimuli are impaired in islets from Aab+ and T1D donors.
- Endothelin-1 action and receptor expression are altered in vascular cells from Aab+ and T1D donors.
- Strong vascular remodeling occurs in the pancreas of Aab+ and T1D donors.

## Introduction

Type 1 diabetes (T1D) is a chronic autoimmune disease that currently has no cure. It is caused by the autoimmune destruction of insulin-producing beta cells in the pancreas, leading to impaired glucose homeostasis and chronic hyperglycemia (1). While the prominent pathological features of T1D pancreata are a marked loss of beta cells and lymphocytic infiltrates in islets [insulitis; (2, 3)], additional pathological abnormalities have been described in endocrine compartments. These relate, in particular, to the structure and function of the islet microvasculature. Not only changes in islet blood vessel diameter, density and composition of the extracellular matrix (ECM) have been reported (4–8), but also functional defects such as increased vascular leakage (9–11) and abnormal islet blood flow dynamics are observed in pre-symptomatic stages of T1D (12, 13). Indeed, studies of individuals at high risk of developing T1D have shown that critical pathogenic mechanisms start before (months to years) clinical diagnosis (3). Because vascular alterations have been detected before clinical symptoms start to develop (“occult” phase), elucidating the specific mechanisms of dysfunction will shed new light into our understanding of T1D pathogenesis.

Vascular networks in pancreatic islets are essential for adequate nutrient sensing by endocrine cells, efficient release of hormones, and timely responses to changes in glycemia (14, 15). Islets are densely vascularized by capillaries made of endothelial cells and covered by pericytes (16, 17). In mice, pericytes provide direct trophic support for beta cells, contributing to their proliferation, maturation and insulin secretion (18–21), or indirectly by participating in ECM synthesis (22). Pericytes may also impact beta cell health and function by maintaining proper blood supply to endocrine cells as they are important regulators of islet capillary diameter and blood flow (17, 23, 24). Importantly, pericyte-mediated changes in islet blood flow are required for proper islet hormone secretion and glucose homeostasis in mice (25). Changes in islet pericyte density, morphology and function occur under pathophysiological conditions. In mouse models of type 2 diabetes, for instance, islet pericytes enlarge and adopt a myofibroblast-like appearance (15, 26, 27), which interferes with their capacity to regulate islet blood flow (24). Although pericytes have important physiological and pathophysiological functions in islets, it is not known what happens to this cell population during T1D development.

Circulating islet autoantibodies (Aabs) are the most robust biomarkers of T1D and are found in 95% of individuals that develop clinical symptoms (28). Single Aab positivity represents the earliest phase in the natural history of islet autoimmunity (2), and antibodies that recognize glutamic acid decarboxylase 65 (GADA) are the most common autoantibody in single Aab+ donors (29). For these reasons, in this study we compared pericyte and vascular responses in islets from Aab-, non-diabetic (control) donors with those in single Aab+ (GADA+) and recent onset T1D organ donors using time-lapse confocal microscopy and living pancreas slices. Our study revealed major changes in the phenotype and function of islet pericytes and, consequently, impaired islet capillary responses to vasoactive stimuli such as high glucose and norepinephrine. We further found abnormal vascular responses to the potent vasoconstrictor endothelin-1 in islets from Aab+ donors, along with altered expression of endothelin-1 receptors in stellate and endothelial cells isolated from the pancreas of Aab+ donors. Our study shows that pericyte dysfunction and impaired islet vasomotion occur at early stages of islet autoimmunity, potentially interfering with the islet response to different metabolic challenges.

## Results

### Pericyte coverage of capillaries is decreased in islets from donors with T1D

Pericytes are crucial for microvascular homeostasis throughout the body (30), but whether they are affected as T1D progresses has not been examined. Here we assessed changes in the density of pericytes in endocrine and exocrine compartments of the human pancreas and compared pericyte to endothelial cell ratio, a major determinant of the tightness of the endothelial barrier (31). We immunostained endothelial cells and pericytes in fixed pancreas slices from organ donors without diabetes (ND), donors with a single islet autoantibody (GADA+; Aab+) and donors with T1D (Figures 1A and S2). Capillaries in islets from ND donors were covered with pericytes that exhibited long cytoplasmic processes that extended along islet endothelial cells (Figure 1B). The average density of pericytes and endothelial cells did not change in islets from Aab+ and T1D donors (Figures 1A and S2), but there was a significant decrease in pericyte to endothelial cell ratio in islets from T1D donors in comparison to ND (Figures 1A,D). Several capillaries in T1D islets lacked pericytes (arrows Figure 1A), or were covered by pericytes that had lost their typical elongated shapes and regular surfaces (Figure 1B). Pericytes in the vicinity of T1D islets also had an abnormal morphology, characterized by multiple cytoplasmic processes or stellate shapes [Figure 1C; (32)], and pericyte coverage of acinar capillaries increased in T1D (Figure 1E).

**Figure 1.**
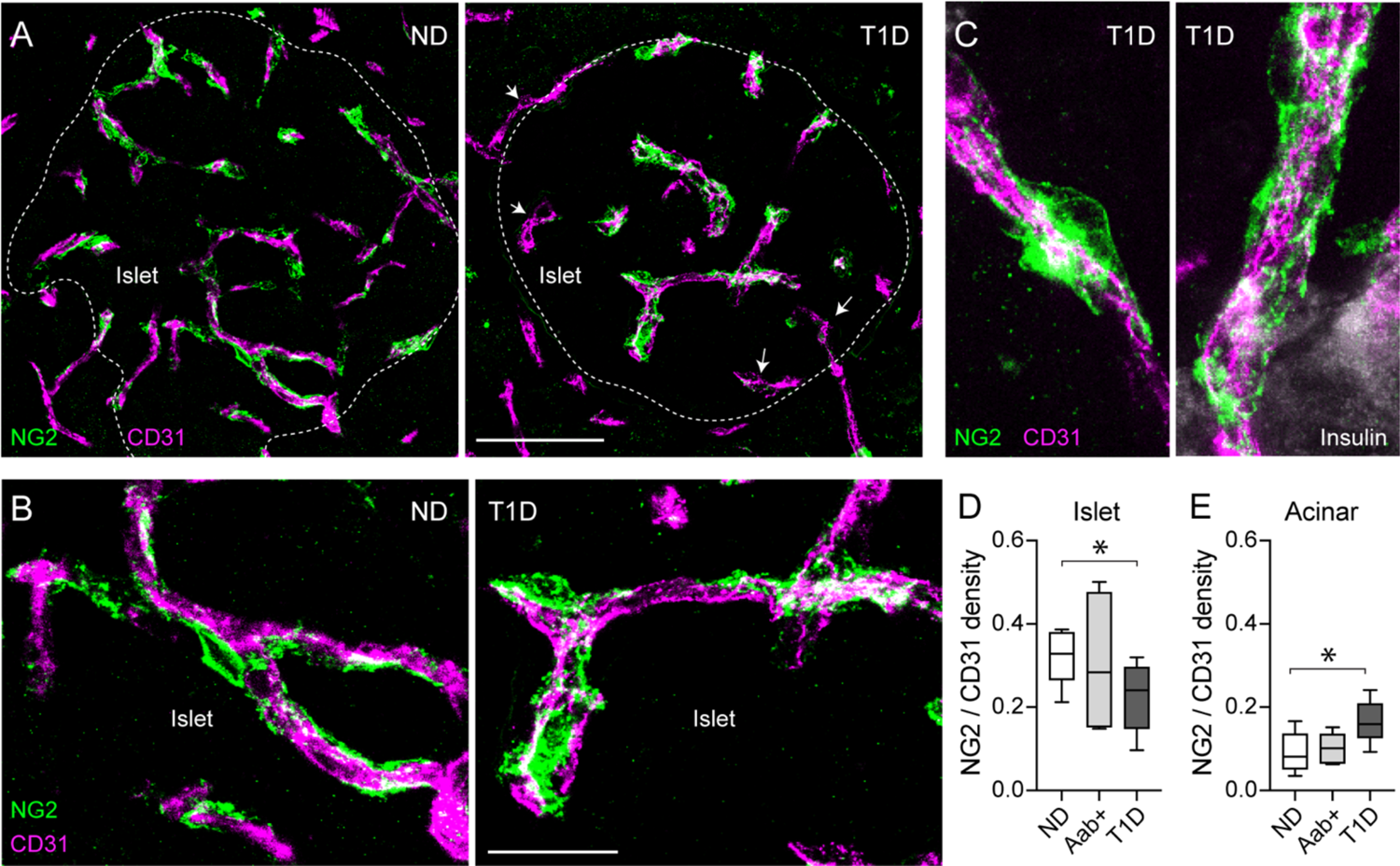
Pericyte coverage of capillaries is decreased in islets from donors with T1D. (A) Maximal projections of confocal images of blood vessels in islets in fixed pancreatic slices from a non-diabetic (ND; nPOD6531) and a T1D donor (T1D duration 1.5y; nPOD6469) immunostained with antibodies against the endothelial cell marker CD31 (magenta) and the pericyte marker neuron-glial antigen 2 (NG2; green). Islets were identified with insulin immunostaining (related to Figure S2; dashed lines indicate islet border). Arrows indicate capillaries in T1D islets lacking pericytes. (B) Projections of confocal images of regions within islets shown in (A). Pericytes with long cytoplasmic processes cover capillaries in ND islets but they have an altered morphology in T1D islets. (C) Projections of confocal images of capillaries at the border of islets in a slice from a T1D donor. Note the altered morphology of the mural cells. (D,E) Estimation of the ratio of pericytes: endothelial cells quantified as NG2 immunstained area: CD31 immunostained area in islets (D) and surrounding acinar tissue (E). N = 9 non-diabetic donors, 7 Aab+ donors (GADA+) and 8 type 1 diabetic donors (T1D). For each donor, around 5-7 islets were imaged and an average value was calculated. * *p* < 0.05 (one-way ANOVA followed by Tukey’s multiple comparisons test). Scale bars = 50 μm (A), 20 μm (B).

**Figure 2.**
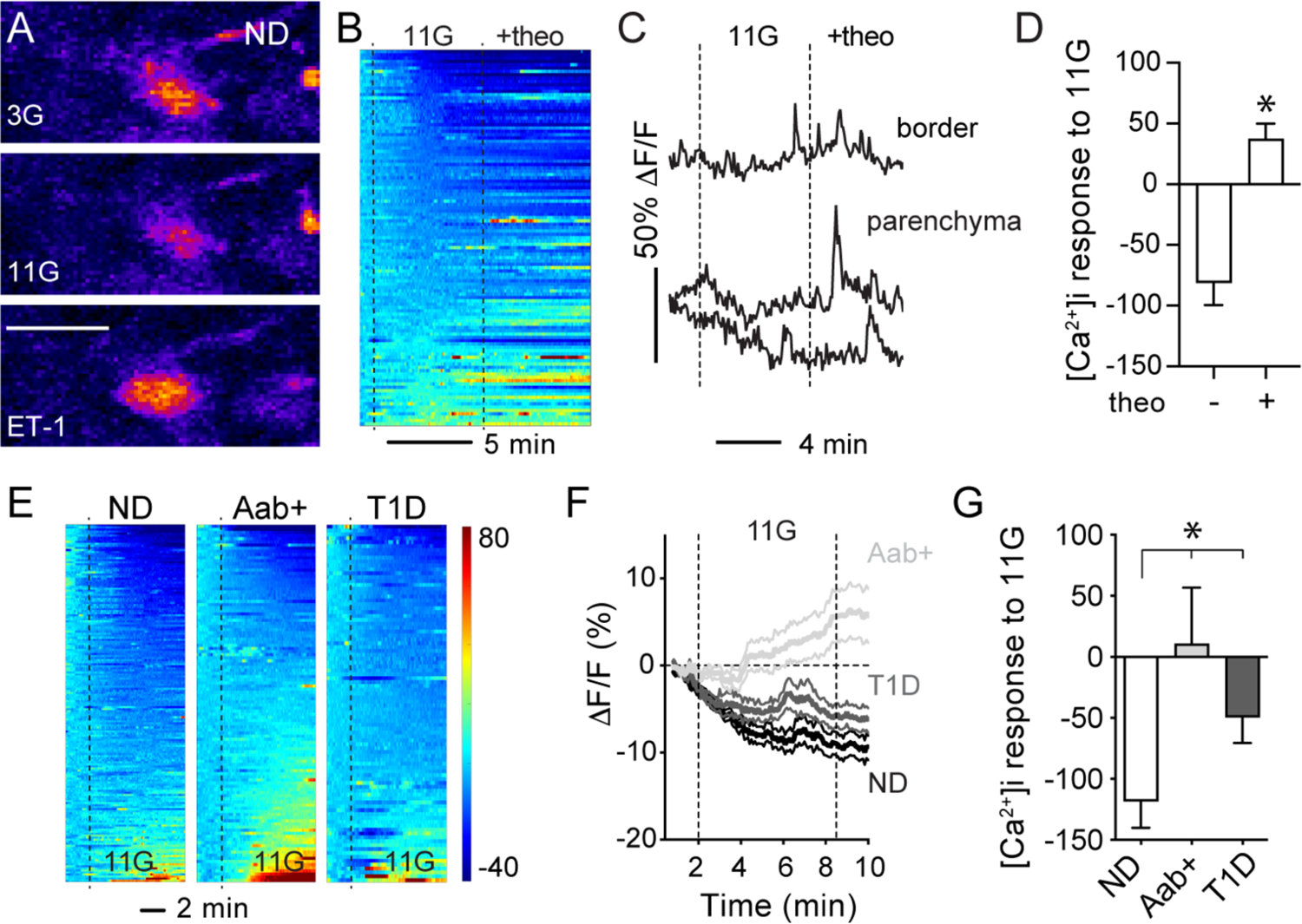
Pericyte calcium responses to glucose are impaired in islets from Aab+ donors. Living human pancreas slices were incubated with the [Ca^2+^]i indicator Fluo4 and with a fluorescent antibody against the pericyte marker NG2 to record changes in pericyte activity (related to Figure S3). (A) Confocal image of a pericyte in an islet from a non-diabetic donor (nPOD6516) in basal glucose concentration (3 mM glucose; 3G), upon stimulation with high glucose (11 mM glucose; 11G) or with endothelin-1 (10 nM; ET-1). Changes in fluo4 fluorescence are shown in a pseudo-color scale. High glucose decreases [Ca^2+^]_i_ in islet pericytes, while ET-1 increases [Ca^2+^]_i_. Scale bar = 10 μm. (B) Heatmap showing changes in fluo4 fluorescence (*Δ*F/F; %) in regions of interest (ROIs) placed around pericytes in islets from non-diabetic (ND) donors elicited by switching extracellular glucose from 3 mM to 11 mM glucose and then applying the non-specific adenosine receptor antagonist theophylline (20 μM) in 11 mM glucose. Each line in the heatmap is a different pericyte. (C) Representative traces showing changes in fluo4 fluorescence (*Δ*F/F; %) elicited by high glucose (11G) alone or in the presence of theophylline (+theo) in individual islet pericytes located at the islet border or within the parenchyma. (D) Quantification of the net area under the curve (AUC) of fluorescence traces as shown in (C). Baseline was defined as the first 10 values in the traces (in 3G or 11G for “theo”). * indicates *p* < 0.05 (paired t-test; N=113 pericytes/5 ND donors). Theophylline reverses the inhibitory effects of high glucose on islet pericytes. (E) Heatmaps showing changes in fluo4 fluorescence (*Δ*F/F; %) elicited by increasing glucose concentration from 3 mM to 11 mM (11G) in pericytes in islets from ND donors (n = 175 pericytes/9 donors), Aab+ (n = 136 pericytes/7 donors) and T1D donors (n = 92 pericytes/4 donors). (F) Traces showing changes in fluo4 fluorescence (*Δ*F/F; %) elicited by high glucose (11G) in islet pericytes. Average (thicker lines) ± SEM values are shown. (G) Quantification of the net AUC of fluorescence traces during stimulation with 11G. * indicates *p* < 0.05 compared to ND (one-way ANOVA followed by Tukey’s multiple comparisons test).

Within this cohort of T1D donors with relatively short disease duration (first four years after diagnosis), we did not observe any significant correlation between the density of islet pericytes and either the duration of diabetes, plasma levels of glycated hemoglobin or the number of circulating antibodies against islet antigens (Figure S2). In summary, our data show that the pericyte to endothelial cell ratio decreases in individuals recently diagnosed with T1D, although pericyte density is similar in islets from single Aab+ and ND donors. Because remaining pericytes in T1D islets exhibited an abnormal morphology, we then asked whether their function was preserved.

### Pericyte calcium responses to glucose are impaired in islets from Aab+ donors

We have previously shown that islet pericytes are very responsive to changes in extracellular glucose concentration (17). In particular, increasing glucose concentration in living mouse pancreas slices decreased cytosolic calcium levels ([Ca^2+^]i) in pericytes and dilated islet capillaries (17). We had further shown that adenosine, most likely produced from ATP co-released with insulin from dense-core granules (33, 34), partially mediated the inhibitory effects of glucose on islet pericytes (17). To examine pericyte responses to high glucose in human islets, we used living human pancreas slices as previously described (23, 25). In this platform, similarly to mouse slices, pericytes remain in their niches within islets and can be visualized with a fluorescent antibody against the pericyte marker and surface proteoglycan neuron-glial antigen 2 [NG2; Figure S3].

Slices were incubated with a membrane permeant calcium indicator (Fluo4) to record changes in [Ca^2+^]i in islet pericytes induced by increasing glucose concentration from 3 mM (3G) to 11 mM (11G). High glucose decreased [Ca^2+^]i in the majority of pericytes in islets from ND donors (Figures 2A,B). As a positive control, we applied the potent vasoconstrictor endothelin-1 which increased [Ca^2+^]i in around ∼50% of the pericyte population in islets from ND donors (Figures 2A and 5). After ∼7 min of being stimulated with high glucose, [Ca^2+^]i of islet pericytes had decreased by 10% of baseline values (Figure 2F). To assess whether adenosine mediated this inhibitory effect of glucose in human islet pericytes, we applied theophylline, a non-specific adenosine receptor antagonist, in the presence of 11 mM glucose. Theophylline (20 μM) increased [Ca^2+^]i in a subset of pericytes in islets from ND donors (Figures 2B,C), reversing the inhibitory effects of high glucose on islet pericytes (Figure 2D). Pericytes activated by theophylline mostly located within the islet parenchyma (Figure 2C).

We then examined pericyte [Ca^2+^]i responses to high glucose in islets from Aab+ and T1D donors. In Aab+ donors, while a subset of islet pericytes were still inhibited by high glucose stimulation (Figure 2E), the majority was activated by high glucose (Figures 2E-G). Glucose led to an average increase of around 5% of [Ca^2+^]i levels in islet pericytes in Aab+ donors (Figure 2F). In contrast, the average effect of high glucose on [Ca^2+^]i levels in pericytes in T1D islets was still inhibitory (Figures 2E-G) but smaller in its magnitude (*Δ*[Ca^2+^]i was only of 5% of baseline values; Figures 2F,G). In summary, our data show that islet pericyte responses to high glucose stimulation are impaired at early stages of T1D. In future studies we will investigate whether endogenous production of adenosine is compromised in islets from Aab+ and T1D donors, potentially interfering with pericyte responses to elevated glucose.

### Vasodilatory responses to glucose are impaired in islets from Aab+ and T1D donors

Because pericytes control islet capillary diameter (17, 23, 25), we then examined capillary responses (vasomotion) induced by high glucose in slices from donors at different stages of T1D. Living pancreas slices were incubated with a fluorescent lectin (*Lycopersicon esculentum* lectin 647) to visualize the islet microvasculature as previously described (23). In most lectin labeled capillaries, a lumen was visible and the average islet vessel diameter was around 5 μm in islets from ND and T1D donors (average basal capillary diameter of 5.0 ± 0.2 for ND donors and 5.7 ± 0.3 for T1D donors; *p*=ns). Interestingly, in islets from Aab+ donors the average islet capillary diameter was slightly reduced under basal, non-stimulatory conditions [average basal capillary diameter of 4.1 ± 0.4 for Aab+ donors; *p*<0.05 between Aab+ and both ND and T1D donors (one-way ANOVA followed by Tukey’s multiple comparisons test)].

Glucose inhibited pericytes in islets from ND donors (Figure 2). This glucose-dependent decrease in [Ca^2+^]i in islet pericytes was associated with a significant increase in capillary diameter in islets from ND donors (Figure 3). Although only a subset (∼60%) of islet capillaries dilated upon glucose stimulation (Figure 3B), high glucose led to ∼5% average increase in islet capillary diameter (Figure 3D). Importantly, vasodilatory responses to glucose were impaired in islets from Aab+ and T1D donors (Figures 3B-D). In summary, changes in [Ca^2+^]i in pericytes triggered by glucose (Figure 2) are not coupled to significant changes in capillary diameter in islets from Aab+ and T1D donors (Figure 3). Islet blood vessels are thus dysfunctional at early stages of T1D and do not dilate upon high glucose stimulation. Impaired vasodilatory responses to glucose can have metabolic consequences as in mice these changes are needed for proper islet hormone secretion upon a glucose challenge (25).

**Figure 3.**
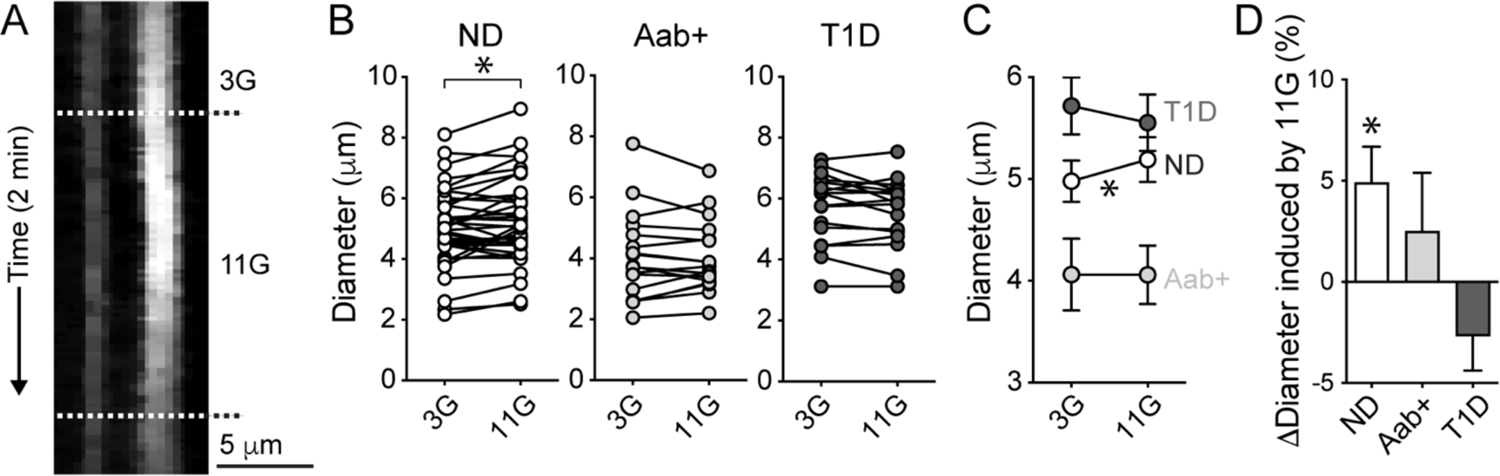
Vasodilatory responses to glucose are impaired in islets from Aab+ and T1D donors. Living human pancreas slices were incubated with a fluorescent lectin from *Lycopersicon esculentum* to label the islet microvasculauture as previously described (23). (A) Temporal projection (reslice image; see Methods) showing changes in diameter of a capillary in an islet from a non-diabetic donor (nPOD6555) induced by switching glucose from 3 mM to 11 mM. (B,C) Quantification of lectin-labeled capillary diameters in living slices from ND (n = 37 capillaries/5 donors), Aab+ (n = 17 capillaries/3 donors) and T1D donors (n = 17 capillaries/5 donors). Mean ± SEM diameter values in 3G and 11G are shown in (C). * indicates *p* < 0.05 (paired t-test comparing values in 3G with 11G for each group of donors). (D) Quantification of the relative changes in capillary diameter (*Δ*Diameter; % baseline) induced by 11G for each group of donors. * indicates *p* < 0.05 (one-sample t-test, theoretical mean = 0).

### Islet microvascular responses to norepinephrine are impaired in Aab+ and T1D donors

Pericytes in islets are innervated by sympathetic nerve fibers and respond to different sympathetic agonists *ex vivo* and *in vivo* (17, 23, 25). Therefore, we then examined whether changes in sympathetic input could have contributed to islet pericyte dysfunction. We first assessed sympathetic innervation patterns in islets at different stages of T1D. To visualize sympathetic nerve fibers, we immunostained tissue slices with an antibody against the sympathetic nerve marker tyrosine hydroxylase (TH). Sympathetic axons could be seen in both endocrine and exocrine compartments of the pancreas of ND, Aab+ and T1D organ donors (Figure S4). The density of TH-positive fibers in islets did not change during these stages of T1D (Figures 4A,B), in contrast with a previous study reporting loss of sympathetic nerves in T1D islets (35). Around 25% of pericytes in islets from non-diabetic individuals were in close contact with sympathetic nerve fibers (Figure 4A). Similarly, sympathetic axons that reached islets in Aab+ and T1D donors also innervated pericytes and the fraction that was innervated remained around 17% and 18%, respectively (Figures 4A and S4).

**Figure 4.**
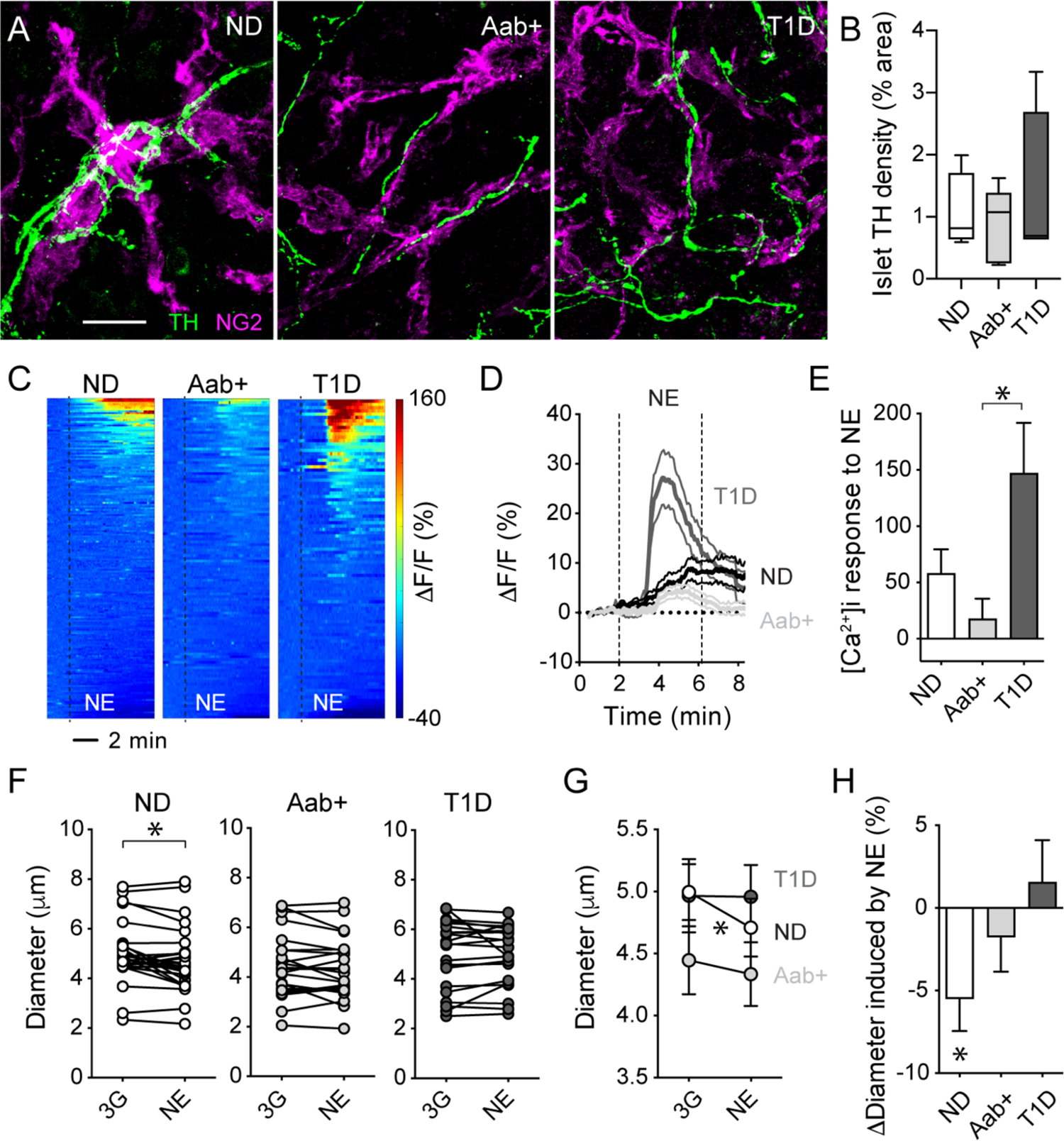
Islet microvascular responses to norepinephrine are impaired in Aab+ and T1D donors. (A) Projections of confocal images of fixed pancreas slices from a non-diabetic (ND; nPOD6546), an Aab+ donor (nPOD6569) and a T1D donor (T1D duration of 2 years; nPOD6566) immunostained for the sympathetic nerve marker tyrosine hydroxylase (TH; green) and the pericyte marker NG2 (magenta). Islets were found using insulin or somatostatin immunostainings (related to Figure S4). Sympathetic nerves reach islets at different stages of T1D and contact a subset of islet pericytes (∼20% of NG2-expressing pericytes). Scale bar = 20 μm. (B) Quantification of the % of islet area immunostained for TH in fixed pancreas slices from non-diabetic (ND), Aab+ and T1D donors. N = 4-5 donors for each group, around 5-7 islets were imaged for each donor and an average value was calculated. (C) Heatmaps showing changes in Fluo4 fluorescence (*Δ*F/F; %) elicited by norepinephrine (NE; 20 μM) in ROIs placed around islet pericytes in living pancreas slices from ND, Aab+ and T1D donors. Each line in each heatmap is a different pericyte. (D) Traces showing changes in fluo4 fluorescence (*Δ*F/F; %) of islet pericytes elicited by NE. Baseline was defined as the first 10 values in fluorescence traces (in 3G). Average (thicker lines) ± SEM values are shown (for ND, n = 182 pericytes/9 donors; for Aab+, n = 117 pericytes/7 donors; for T1D, n = 97 pericytes/5 donors). (E) Quantification of the net AUC of individual fluorescence traces as those shown in (D). * indicates *p* < 0.05 (one-way ANOVA followed by Tukey’s multiple comparisons test). (F,G) Quantification of changes in diameter of islet capillaries induced by norepinephrine (NE; 20 μM; applied in 3G) in slices from ND (n=29 capillaries/7 donors), Aab+ (n=23 capillaries/5 donors) and T1D donors (n=22 capillaries/5 donors). Individual diameter values are shown in (F), while mean ± SEM diameter values in 3G and NE are shown in (G). * indicates *p* < 0.05 (paired t-test comparing values in 3G with NE for all the groups). (H) Quantification of the relative changes in capillary diameter (*Δ*Diameter; % baseline) induced by NE for each group of donors. * indicates *p* < 0.05 (one-sample t-test, theoretical mean = 0).

We then examined islet microvascular responses to the sympathetic agonist norepinephrine by confocal imaging of living pancreas slices. As previously shown (23), norepinephrine (20 μM) increased [Ca^2+^]i levels in a subset of pericytes in islets from ND donors (Figure 4C). Indeed, only around 30% of islet pericytes were activated by norepinephrine, and we have previously shown that responding pericytes were those innervated by sympathetic nerve fibers (25). Increases in pericyte [Ca^2+^]i induced by norepinephrine were accompanied by constriction of a subset of islet capillaries (Figures 4F-H). Norepinephrine induced a ∼5% decrease in islet capillary diameter in ND donors (Figure 4H). Importantly, pericyte [Ca^2+^]i responses to norepinephrine were greatly diminished in islets from Aab+ donors (Figures 4C-E) and norepinephrine stimulation did not constrict islet capillaries (Figures 4F-H). In contrast, pericytes in islets from T1D were very responsive to norepinephrine and this agonist induced a robust increase in pericyte [Ca^2+^]i in islets from T1D donors (Figures 4C-E). However, norepinephrine stimulation failed to elicit vasoconstriction of capillaries in islets from T1D donors (Figures 4F-H). Our data again show that changes in [Ca^2+^]i in islet pericytes are uncoupled from efficient vasodilation/vasoconstriction of the microvasculure in T1D islets. In particular, our data suggest that islet pericytes lose their capacity to respond to sympathetic nervous input at early stages of T1D. We had previously shown that *in vivo* activation of pericytes with a sympathetic agonist constricts islet capillaries and decreases blood flow in intraocular islet grafts, which decreases plasma insulin levels (25). Abnormal pericyte/microvascular responses to sympathetic agonists in human islets may compromise sympathetic-induced changes in blood flow and impact islet hormone secretion at early stages of the disease.

### Altered endothelin-1 vascular effects and receptor expression in Aab+ and T1D islets

Pericytes in mouse and human islets are also very responsive to endothelin-1 [Figure 2A; (17, 23)], an extremely potent vasoconstrictor peptide that is produced by endothelial cells throughout the body (36). Indeed, exogenous endothelin-1 (10 nM) increased [Ca^2+^]i levels specifically in islet pericytes (Figure 5A), and decreased capillary diameter by ∼8% in islets from ND donors (Figures 5D,E). Pericyte [Ca^2+^]i responses to endothelin-1 were diminished in islets from Aab+ donors (Figures 5B,C), and endothelin-1 administration did not result in significant vasoconstriction (Figures 5D,E). Interestingly, pericytes in islets from Aab+ donors were still very responsive to another potent vasoconstrictor peptide, angiontensin II (Figure 5F). Similarly to their “preserved” [Ca^2+^]i responses to high glucose or norepinephrine, pericytes in T1D islets were potently activated by endothelin-1 (Figures 5A-C), but again [Ca^2+^]i responses were not associated with significant changes in capillary diameter (Figures 5D,E).

**Figure 5.**
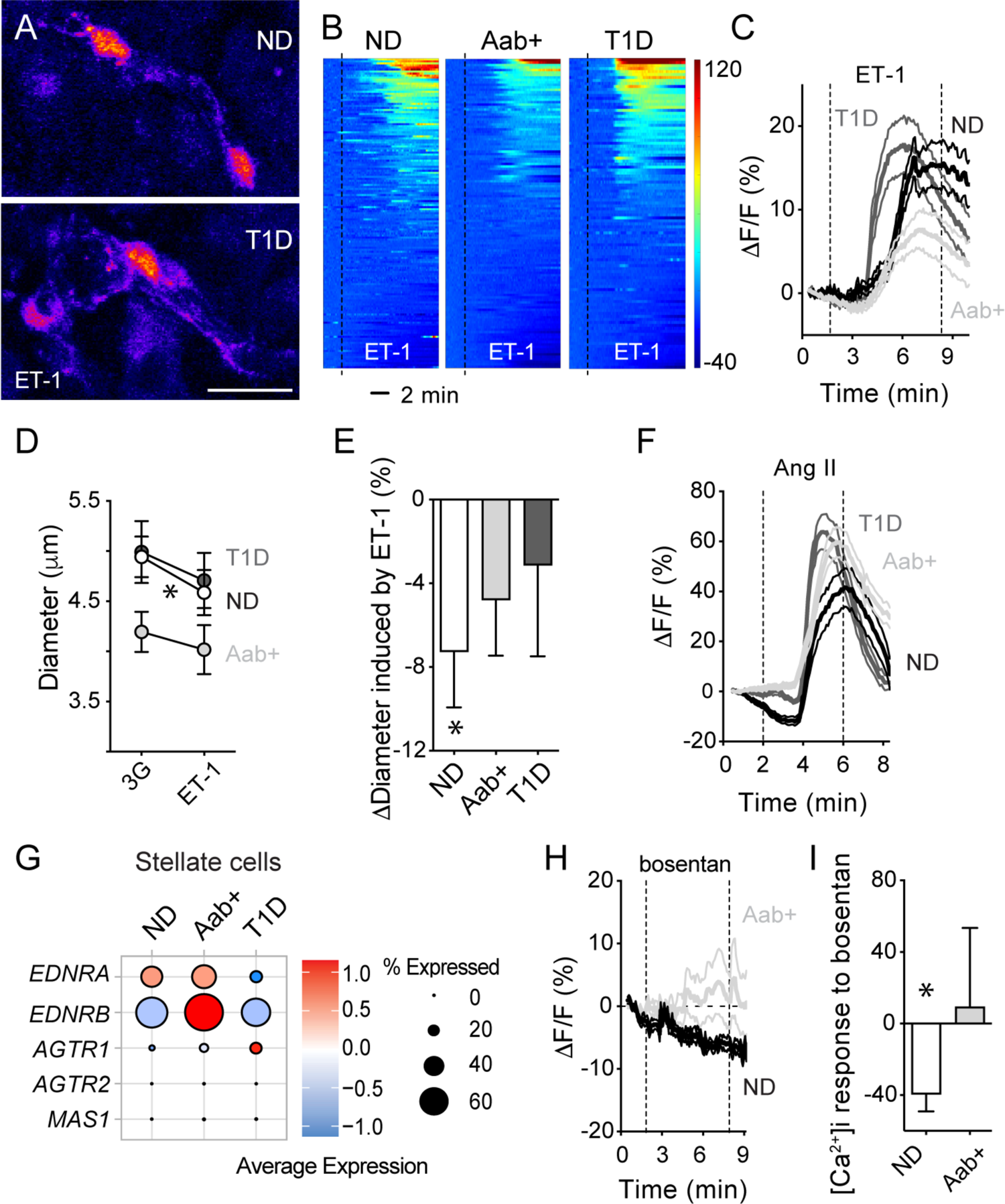
Altered endothelin-1 vascular effects and receptor expression in Aab+ and T1D islets. (A) Islet pericytes in living pancreas slices from a ND donor (nPOD6516) and a T1D donor (T1D duration 0.6 years; nPOD6551) responding to ET-1 (10 nM) are shown in a pseudo-color scale. Pericytes were identified with anti-NG2-alexa647 antibody as previously described (23). Note the different shape of islet pericytes in ND and T1D slices. Scale bar = 20 μm. (B) Heatmaps showing changes in fluo4 fluorescence (*Δ*F/F; %) elicited by endothelin-1 (ET-1) in ROIs placed around islet pericytes in slices from ND, Aab+ and T1D donors. Each line in each heatmap is a different pericyte. (C) Traces showing changes in fluo4 fluorescence (*Δ*F/F; %) of islet pericytes elicited by endothelin-1 (ET-1). Baseline was defined as the first 10 values in fluorescence traces (in 3G). Average (thicker lines) ± SEM values are shown (for ND, n = 166 pericytes/9 donors; for Aab+, n = 125 pericytes/7 donors; for T1D, n = 104 pericytes/5 donors). (D) Quantification of changes in diameter of islet capillaries induced by endothelin-1 (ET-1; 10 nM; applied in 3G) in slices from ND (n=32 capillaries/6 donors), Aab+ (n=21 capillaries/5 donors) and T1D donors (n=21 capillaries/5 donors). Mean ± SEM diameter values in 3G and ET-1 are shown. * indicates *p* < 0.05 (paired t-test comparing values in 3G with ET-1 for all the groups). (E) Quantification of the relative changes in capillary diameter (*Δ*Diameter; % baseline) induced by ET-1 for each group of donors. * indicates *p* < 0.05 (one-sample t-test, theoretical mean = 0). (F) Traces showing changes in fluo4 fluorescence (*Δ*F/F; %) of islet pericytes elicited by angiotensin II (Ang II; 100 nM). Baseline was defined as the first 10 values in fluorescence traces (in 3G). Average (thicker lines) ± SEM values are shown (for ND, n = 91 pericytes/3 donors; for Aab+, n = 138 pericytes/5 donors; for T1D, n = 71 pericytes/3 donors). (G) Dotplots showing average expression levels of genes encoding endothelin receptors (*EDNRA* and *EDNRB*) and angiotensin II receptors (*AGTR1*, *AGTR2* and *MAS1*) in stellate cells isolated from ND, Aab+ or T1D pancreata. Data were extracted from HPAP database [(37); see Methods and Figure S5. (H) Traces showing changes in fluo4 fluorescence (*Δ*F/F; %) of islet pericytes elicited by the dual endothelin receptor antagonist bosentan (100 nM in 3G). Baseline was defined as the first 10 values in fluorescence traces (in 3G). Average (thicker lines) ± SEM values are shown (for ND, n = 25 pericytes/3 donors; for Aab+, n = 34 pericytes/3 donors). (I) Quantification of the net area under the curve (AUC) of fluorescence traces as those shown in (H). * indicates *p* < 0.05 (one-sample t-test, theoretical mean = 0).

Using a publicly available database containing single cell RNAseq data from different cells in the human pancreas [from the Human Pancreas Analysis Program; (37)], we analyzed potential changes in gene expression of endothelin-1 receptors in stellate cell and endothelial cell clusters at different stages of T1D. Stellate cell clusters in the pancreas include pericytes evidenced by the expression of pericyte identity genes (e.g. *CSPG4*, *PDGFRB* and *ACTA2*) by cells within this cluster [Figure S5; (38)]. Endothelin-1 exerts its effects by binding to two G-protein-coupled receptors - ET_A_ and ET_B_ ^-^ encoded by *EDNRA* and *EDNRB* genes, respectively. Stellate cells in non-diabetic pancreas express mainly *EDNRA* which mediates the vasoconstrictor effects of endothelin-1 [Figure 5G; (39)]. Interestingly, there was a significant upregulation (fold change *vs.* ND = 1.5) of *EDNRB* in stellate cells from Aab+ donors, while *EDNRA* was downregulated in stellate cells from T1D donors (Figure 5G; fold change *vs.* ND = 0.8). There were no significant changes in expression levels of angiotensin II receptors (Figure 5G), adenosine or adrenergic receptors in stellate cells (data not shown). Similarly, *EDNRB* was upregulated in endothelial cells from Aab+ donors (fold change *vs.* ND = 1.3). In line with altered endothelin-1 receptor expression by vascular cells in Aab+ pancreata, antagonizing endothelin-1 receptor signaling with the dual receptor antagonist bosentan (100 nM) affected differentially [Ca^2+^]i levels in pericytes in islets from ND and Aab+ donors (Figures 5H,I). In particular, bosentan significantly decreased pericyte [Ca^2+^]i levels in islets from ND donors in agreement with endothelin-1 exerting its vasoconstrictor effects by activating ET_A_ receptors (39). In contrast, bosentan did not alter pericyte [Ca^2+^]i levels in islets from Aab+ donors, most likely because of the upregulation of ET_B_ receptors in endothelial cells which lead instead to vasodilation (39). In summary, our data show that altered endothelin-1 vascular effects and receptor expression occur in Aab+ and T1D donor pancreata.

### Altered pericyte and stellate cell phenotypes in islets from Aab+ and T1D donors

Given pericytes’ abnormal function and impaired vasomotive responses of capillaries in islets from Aab+ and T1D donors, we hypothesized that islet pericytes were turning into ECM producing myofibroblasts and losing their capacity to control islet capillary diameter. Indeed, conversion of pericytes into myofibroblasts had been previously reported in a mouse model of islet fibrosis (40). To test this hypothesis, we examined by immunohistochemistry pericytic expression of the ECM protein collagen type IV. The density of this basement membrane component increased significantly in islets from T1D donors (Figures 6A,B). Islet pericytes were found embedded within these collagen type IV networks (Figure 6C). Importantly, the proportion of colocalization of the pericyte marker NG2 with collagen IV increased significantly in islets from Aab+ and T1D donors (Figure 6D), indicative of an increased contribution of islet pericytes to the synthesis of this ECM protein. We then assessed pericytic expression of myofibroblast markers such as alpha smooth muscle actin (*α*SMA) and periostin (41). Pericytes could be seen in close proximity to periostin aggregrates in islets from Aab+ donors (Figure S6), but this matricellular protein could be detected only in a subset of islets at different stages of T1D (Figure S6). Moreover, the proportion of NG2-expressing pericytes that also expressed *α*SMA inside islets or at the islet border also increased from around 30-40% in ND donors to ∼70% in islets from T1D donors (Figure S6).

**Figure 6.**
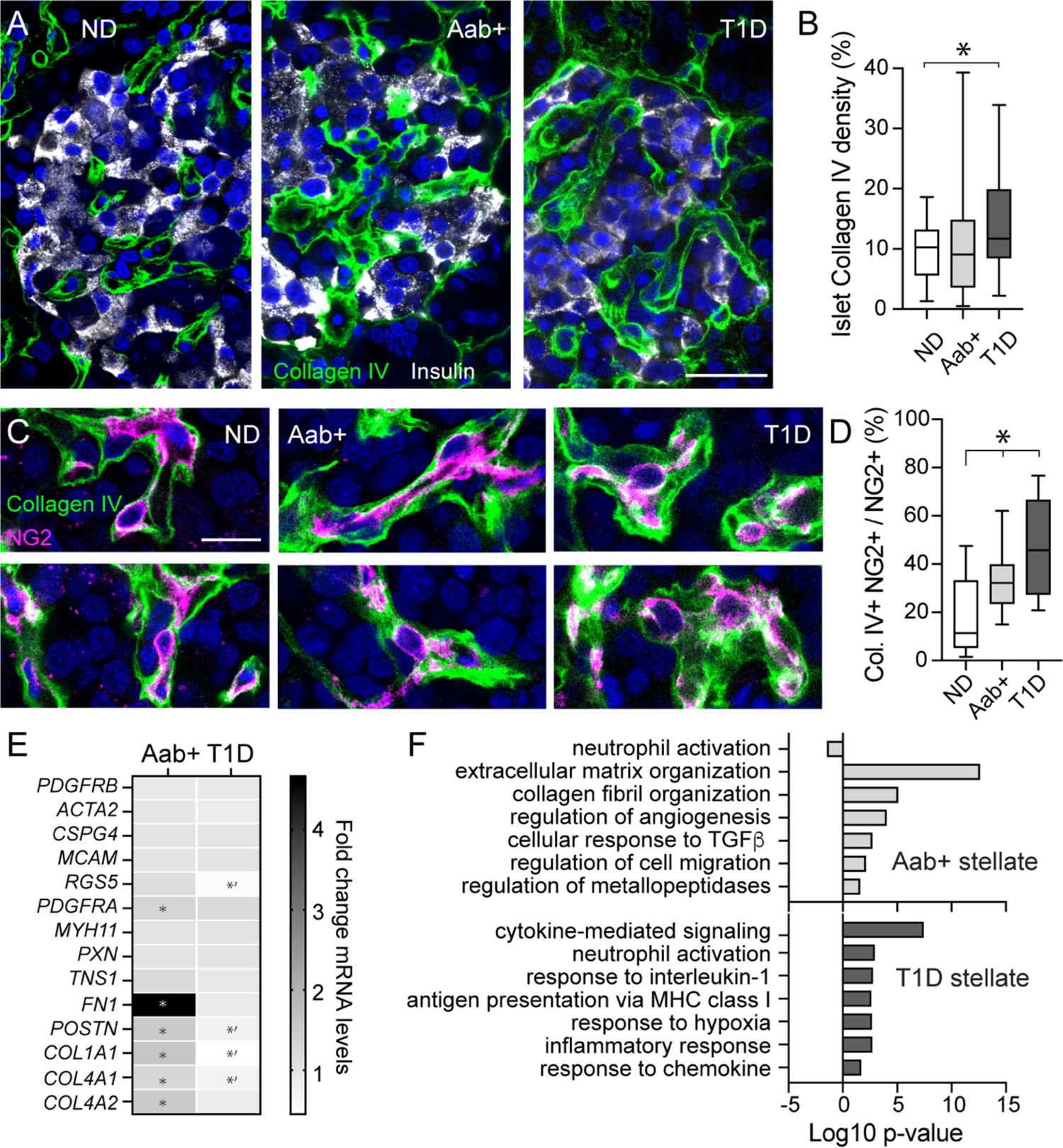
Altered pericyte and stellate cell phenotypes in islets from Aab+ and T1D donors. (A) Confocal images of islets in fixed pancreatic slices from a ND (nPOD6539), a Aab+ (nPOD6573) and a T1D donor (nPOD6563) immunostained with antibodies against insulin (gray) and the basement membrane component collagen type IV (green). (B) Quantification of the % of islet area immunostained with collagen IV in slices from non-diabetic (ND, n = 34 islets/5 donors), GADA+ (Aab+, n = 36 islets/5 donors) and type 1 diabetic donors (T1D, n = 28 islets/3 donors). * indicates *p* < 0.05 (one-way ANOVA followed by Tukey’s multiple comparisons test). (C) Confocal images of NG2-labeled pericytes (magenta) embedded in the basement membrane in islets from ND, Aab+ and T1D donors above, showing colocalization of NG2 and collagen IV (green). Colocalization between NG2 and collagen IV appears in white. (D) Mander’s correlation coefficient estimating colocalization between NG2 and collagen IV in confocal images of islets from ND (n = 19 islets/5 donors), Aab+ (n = 24 islets/4 donors) and T1D donors (n = 17 islets/3 donors). * indicates *p* < 0.05 (one-way ANOVA followed by Tukey’s multiple comparisons test). (E) Change in expression of different genes in stellate cells from Aab+ or T1D donors compared to non-diabetics. Data are extracted from HPAP database. Included are genes expressed by pericytes (*PDGFRB, CSPG4, MCAM, RGS5, ACTA2*), fibroblasts/myofibroblasts (*PDGFRA*, *COLA1A1, COL4A1, COL4A2, FN1, MYH11, PXN, TNS1* and *POSTN*). * indicates significant upregulation (fold change (FC) > 1.2) while *’ indicates significant downregulation (FC < 0.8). (F) Gene ontology analysis of the top 6 biological processes that are significantly upregulated in stellate cells from Aab+ (light gray) or T1D pancreata (dark gray) in comparison to stellate cells from non-diabetics. While stellate cells from Aab+ donors are mostly involved in ECM organization and vascular remodeling, they acquire more immunological functions in T1D pancreata. Scale bars = 20 μm (A), 10 μm (C).

To examine in more detail changes in the phenotype of pericyte/stellate cells in the pancreas of Aab+ and T1D donors, we used the HPAP scRNAseq database. We analyzed the expression of different pericyte identity genes (*PDGFRB, CSPG4, MCAM, RGS5, ACTA2*), genes associated with a myofibroblast phenotype such as smooth muscle myosin heavy chain (*MYH11*), periostin (*POSTN*), genes encoding focal adhesion proteins such as paxillin (*PXN*), tensin (*TSN1*) and ECM proteins fibronectin (*FN1*) and collagens (e.g. *COL1A1, COL4A1, COL4A2*) in stellate cell populations from non-diabetics, Aab+ and T1D donors (Figure 6E). There were no significant transcriptional alterations of most pericyte genes except a significant downregulation of *RGS5* in stellate cells from T1D donors (Figure 6E). *RGS5* (regulator of G-protein signaling 5) is a pericyte-specific gene whose expression is reduced in vessels with poor pericyte coverage compared to a mature vasculature (42). Genes encoding fibronectin, periostin and collagen subunits were significantly upregulated in stellate cells from Aab+ donors, but downregulated in T1D stellate cells when compared to levels in non-diabetics (Figure 6E). Moreover, there was an interesting switch in gene programs significantly upregulated in stellate cells in Aab+ and in T1D donors: while stellate cells in Aab+ pancreata upregulated pathways involved in ECM production, collagen organization and cell migration, stellate cells in T1D donors exhibited more immunoregulatory functions (Figure 6F). Our data show that are major changes occurring in the phenotype of pericytes and stellate cells in the pancreas at different stages of T1D, that could have impacted their function.

### Strong vascular remodeling occurs in the pancreas of Aab+ and T1D donors

Lastly, we examined transcriptional alterations of pancreatic endocrine, endothelial and stellate cell populations potentially associated with the vasculature. We hypothesized that microvascular dysfunction could have interfered with islet blood perfusion, creating a hypoxic environment and triggering compensatory vascular and metabolic responses. Using the HPAP database, we looked for changes in the expression of genes known to be involved in tissue responses to hypoxia [e.g. during cancer; (43, 44)]. These include genes encoding hypoxia-inducible transcription factors (e.g. *HIF1A, HIF3A, ARNT*) or proteins involved in HIF1*α* stabilization/degradation (e.g. *VHL*), genes involved in angiogenesis and vascular remodeling such as adrenomedullin (*ADM*) and different angiogenic factors (e.g. *ANGPTL4, VEGFA*) and their receptors (e.g. *NRP1, KDR, FLT1*), genes involved in cell migration (e.g. *ENPP2*, *MIF*), glucose transport (*SLC2A1, SLC2A3*) and metabolism [e.g. phosphofructokinase (*PFKP*), pyruvate dehydrogenase kinase isozyme 4 (*PDK4*), lactate dehydrogenase A (*LDHA*) and phosphoglycerate kinase (*PGK1*)]. We noticed that the gene encoding the vasodilator peptide adrenomedullin (*ADM*) was upregulated in different cell types in Aab+ and T1D pancreata (Figure 7A). In addition, there was a significant upregulation of genes encoding different angiogenic and growth factors and their respective receptors in pancreatic endothelial and stellate cells in Aab+ donors (Figure 7A), indicating potential active vascular remodeling. Vascular alterations were accompanied by metabolic changes indicated by significant upregulation of *PDK4* and *LDHA* in endocrine alpha and beta cells in T1D pancreata. These data show that transcriptional alterations related to the vasculature are present in the pancreas of single Aab+ donors, similarly to what had been described for T1D (45).

**Figure 7.**
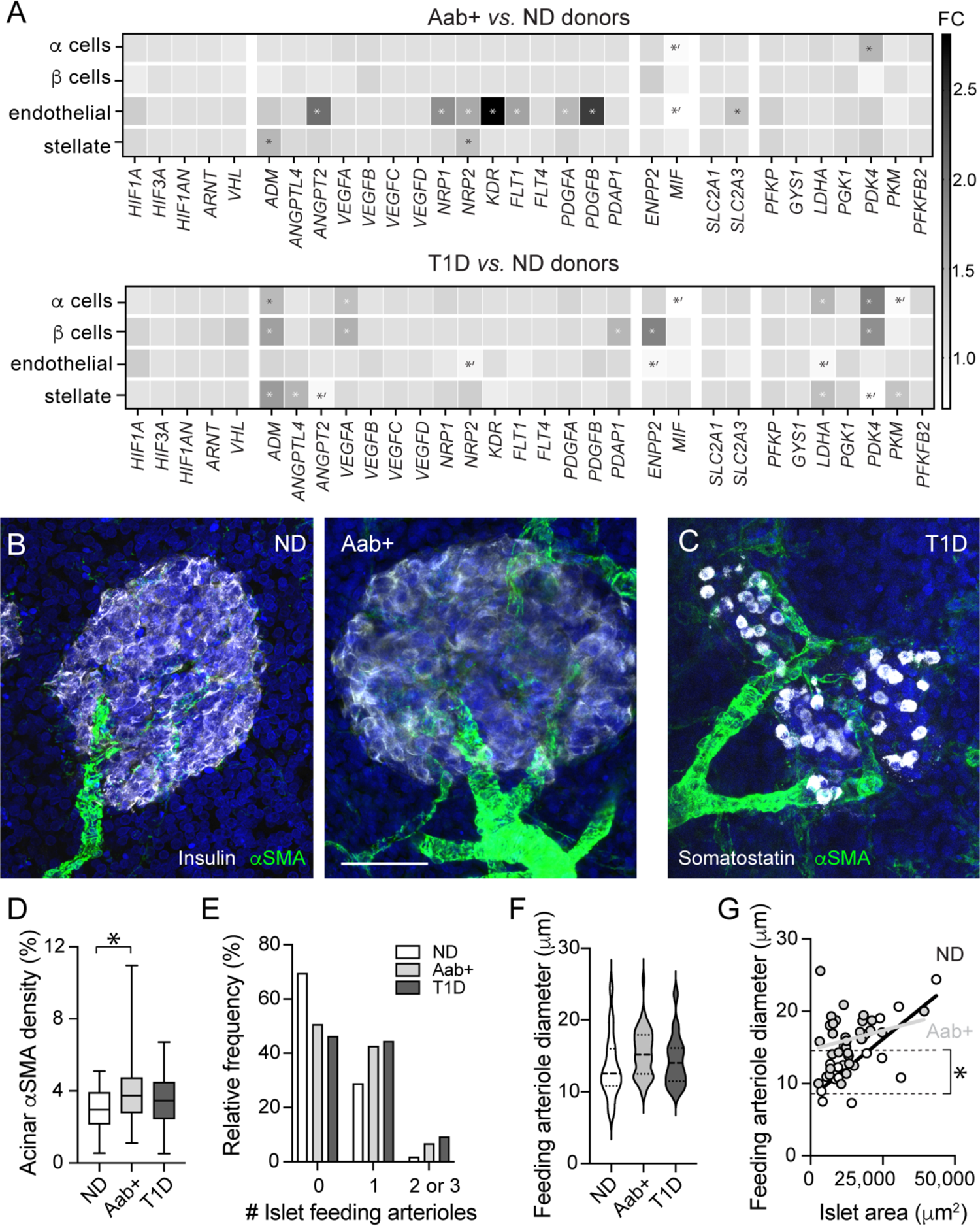
Vascular remodeling occurs in Aab+ and T1D donor pancreata. (A) Changes in the expression of different genes potentially associated with hemodynamic alterations in endocrine alpha and beta cells, endothelial and stellate cells from non-diabetic (ND), Aab+ and T1D donor pancreata. Data have been extracted from the HPAP database. Shown are fold change (FC) in expression levels of genes involved in a first response to hypoxia (*HIF1A, HIF3A, HIF1AN, ARNT, VHL*), genes involved in vasodilation (*ADM*), angiogenesis and vascular remodeling (*ANGPTL4, ANGPT2, VEGFA-D, NRP1-2, KDR, FLT1, FLT4, PDGFA, PDGFB, PDAP1*), cell migration (*ENPP2, MIF*), and metabolism (*SLC2A1, SLC2A3, PFKP, GYS1, LDHA, PGK1, PDK4, PKM, PFKFB2*) in different cell populations in Aab+ (top) or T1D pancreata (bottom) compared to expression levels in non-diabetics. * indicates significant upregulation (FC > 1.2), *’ indicates significant downregulation (FC < 0.8; see Methods). (B,C) Maximal projections of confocal images of islets in fixed pancreatic slices from a ND donor (left; nPOD6539), a Aab+ donor (middle; nPOD6558) and a T1D donor (T1D duration 2.5y; nPOD6480) immunostained for *α*SMA (green) and insulin (gray; ND and Aab+ donors; B) or somatostatin (gray; T1D donor; C). Multiple feeding arterioles irrigate islets in Aab+ and T1D pancreases. Scale bar = 40 μm. (D) Quantification of the % of exocrine tissue area immunostained with *α*SMA in pancreas slices from ND (n = 39 islets/5 donors), Aab+ (n = 64 islets/6 donors) and T1D donors (n = 52 islets/7 donors). * indicates *p* < 0.05 (one-way ANOVA followed by Tukey’s multiple comparisons test). (E) Histogram showing the proportion of islets (in %) containing none, a single (1) or multiple (2–3) feeding arterioles in ND, Aab+ and T1D pancreata. (F) Quantification of the diameter of feeding arterioles in islets from non-diabetics (ND, n = 20 islets/7 donors), Aab+ (n = 40 islets/6 donors) and T1D donors (n = 35 islets/6 donors). (G) Correlation between the islet area and the diameter of the corresponding feeding arteriole. In non-diabetic donors (black symbols and line), there is a linear relationship between these two parameters (R^2^=0.61, *p*=0.0001), while in Aab+ donors (gray symbols and line), this dependency disappears (R^2^=0.05, *p*=0.23). In addition, Y-intercept values (elevations) of the two linear regressions are significantly different (*p*=0.0013).

To visualize whether vascular remodeling occurred in the pancreas, we immunostained pancreatic tissue from Aab+ and T1D donors with an antibody against alpha-smooth muscle actin (*α*SMA). *α*SMA is a contractile protein expressed by mural cells - pericytes and smooth muscle cells – and myofibroblasts [Figure S6; (17, 40)]. Smooth muscle cells can be easily distinguished from other cell types given their circular cytoplasmic processes and multiple rings they form around arterioles or arteries (Figures 7B,C). We observed an increase in the number of arterioles irrigating islets in Aab+ and T1D pancreata (Figures 7B-D). Indeed, while only around ∼30% of islets in non-diabetic donors had one feeding arteriole, around 50% of islets in Aab+ and T1D pancreata had one or more than one feeding arteriole (Figure 7E). There was no significant difference in the diameter of islet feeding arterioles (Figure 7F). Interestingly, while in islets from ND donors there was a significant correlation between the islet area and the diameter of the corresponding feeding arteriole (R^2^=0.6, *p*=0.0001), this correlation was lost in islets from Aab+ donors (R^2^=0.05, *p*=ns; Figure 7G). In T1D donors, this dependency was reestablished ((R^2^=0.2, *p*=0.01). Interestingly, Y intercept values (elevations) were significantly higher for both Aab+ and T1D donors, suggesting that Aab+ and T1D islets may be equipped to receive more blood than what would be needed to supply those tissue areas (Figure 7G). This apparent increase in vascular supply of Aab+ and T1D islets could have occurred as a compensatory responses to impaired vascular perfusion of endocrine tissues at these stages.

## Discussion

In this study, we inspected islet pericytes and compared their density, phenotype, response profile and gene expression in non-diabetic donors with those in individuals at different stages of T1D: single Aab+ (GADA+) and T1D donors with relatively short disease duration (from 0-4 years). We found that, while there are only modest changes in the density of these vascular cells, their function and phenotype are significantly impaired at early stages of the disease. Islet pericytes are needed for proper insulin secretion and glucose homeostasis in mice (25, 46). Their dysfunction could compromise insulin release from endocrine beta cells, exacerbating beta cell stress and potential failure. In this study we have used living pancreas slices to monitor islet pericyte and capillary responses. Although this is an *ex vivo* setting and important regulators of vascular function are missing (input from the central nervous system, blood flow and shear stress, etc), we believe that our approach reflects the *in vivo* situation to a high degree, giving important insights into the pathophysiology of T1D in humans.

A variety of mechanisms are believed to contribute to the pathogenesis of T1D, which could explain the significant heterogeneity and pathological features of this disease (3). T1D is considered a disease of both the immune system and the beta cell, where not only islet-specific autoreactive T cells mistakenly destroy “healthy” beta cells, but also stressed beta cells may trigger an autoimmune attack against themselves (47). A defective islet microvasculature could contribute in both ways, either by facilitating immune cell infiltration into the pancreas, or by compromising vascular perfusion and lowering oxygen levels, exacerbating beta cell stress. Vasculopathy is observed in different tissues throughout the diabetic body, and it is usually considered a consequence of diabetes (48). However, some of the metabolic challenges associated with a diabetic environment (e.g. hyperglycemia, inflammation and oxidative stress) that damage vascular cells (49) may already be present in a prediabetic state in the pancreas [e.g. inflammation in exocrine tissue; (50)]. Indeed, different groups had reported significant anatomical and functional alterations of the islet microvasculature before the onset of symptoms (5, 7, 10, 13, 51, 52), but the cellular and molecular mechanism underlying these defects were missing.

Pericytes are crucial for microvascular homeostasis and stability throughout the body (30, 53). Substantial evidence indicates that pericyte loss (or dysfunction) leads to microvascular instability, disrupting the barrier properties of the endothelium and to the formation of microaneurysms, microhemorrhages, acellular capillaries and capillary nonperfusion (54, 55). In this study we observed a significant decrease in the pericyte to endothelial cell ratio in islets from T1D donors (Figure 1), which is a major determinant of the tightness of the endothelial barrier (31). These data are line with a previous study using magnetic resonance imaging and nanoparticles that revealed abnormal vascular integrity and leukocyte infiltration in the pancreas of T1D patients within six months of diagnosis (51). Besides controlling transendothelial permeability, pericytes have also been shown to actively participate in immunosurveillance in different tissues (56). In response to inflammatory mediators, pericytes express adhesion molecules and chemoattractant factors controlling leukocyte migration *in vivo* (57, 58). Future studies investigating the role of pericytes as active components/mediators of immune responses in islets would be needed to understand the potential impact of their dysfunction in (pre-)diabetes.

Pericytes are also important regulators of capillary blood flow in different tissues such as the brain, the retina and the islet (17, 59–62). Interestingly, there is compelling evidence that islet blood flow is disturbed during conditions of impaired glucose tolerance and overt diabetes, but it is not known if these disturbances are of pathogenic importance (63). Researchers have reported both increases and decreases in islet blood flow in animal models of both type 1 and type 2 diabetes. It seems that at early stages of deranged metabolism blood flow increases in islets but, as the disease progresses, it is then followed by a decrease in islet blood perfusion (64). A recent study using ultrasound contrast imaging revealed altered islet blood flow dynamics in mice before T1D onset, characterized by increased perfusion velocity but reduced perfusion volume (13). These findings in mouse models are in line with our observations in islets from Aab+ and T1D donors, which have a higher number of feeding arterioles (Figure 7) but either decreased islet capillary diameter (as we show in this manuscript) or decreased vessel density as previously shown (8).

In this study, we also found that changes in [Ca^2+^]i in pericytes in islets from Aab+ and T1D donors, elicited either by high glucose, norepinephrine or endothelin-1, are not associated with significant changes in islet vessel diameter (Figures 2-5). We do not know why islet pericytes become dysfunctional. We hypothesized that, given pericytes’ enormous plasticity and postnatal undifferentiated nature, they have switched their phenotype and turned into ECM producing myofibroblasts [activated stellate cells; (Figure 6)]. Indeed, human T1D is characterized by a modified islet ECM (65), and islet pericytes are capable of producing ECM proteins under physiological and pathophysiological conditions (22, 40). A switch in the pericyte phenotype – from an excitable, electrically coupled mural cell to a non-excitable fibroblast-like cell - would have functional consequences. Electrical coupling between mural cells allows the sequential spread of depolarizations to develop synchronous calcium transients within their network (66). Because these rhythmical contractions of mural cells are required for efficient vasomotion (67), the conversion of islet pericytes, even if only a subset of them, into myofibroblasts could produce barriers for conduction of signals, interfering with effective vasodilation or vasoconstriction. Pericyte dysfunction at early stages of the disease would interfere with the islet’s capacity to induce acute changes in its blood flow, triggered for instance upon hyperglycemia or changes in sympathetic tonus (25).

Our study further suggests that altered endothelin-1 signaling/action in the pancreas may contribute to vascular dysfunction. Endothelins are among the most potent vasoconstrictors in the body. Previous studies had shown that endothelin-1 was present in the human pancreas and its expression to be regulated by hypoxia (68, 69). Here we found that pericyte [Ca^2+^]i responses to exogenous endothelin-1 and to a dual endothelin-1 receptor antagonist (bosentan) were altered in islets from Aab+ donors (Figure 5). A potential explanation for these results is that *EDNRB* is significantly upregulated in endothelial cells from Aab+ donors (Figure 5). Upregulation of ET_B_ receptors in islets has been previously observed during hypoxia *in vitro* and *in vivo* in islet grafts early after transplantation when these were still avascular (69). In different tissues throughout the body (e.g. kidney, liver and lungs), ET_B_ receptors have also been shown to play a critical role scavenging pathological levels of endothelin-1 from circulation and targeting it to degradation (70, 71). Upregulation of ET_B_ receptors by vascular cells in the pancreas of Aab+ donors could have been a compensatory response of the tissue to limit the vasoconstrictive effects of endothelin 1 which, besides its vasoactive functions, also stimulates proliferation of fibroblasts and ECM production (39, 68).

To summarize, islets are equipped with vascular networks that not only support endocrine cell health and maturation but also their function as they enable proper glucose sensing and hormone secretion. Our study shows that the islet microvasculature is dysfunctional already in islets from single Aab+ donors. Whether islet vascular dysfunction impairs *in vivo* endocrine cell responses to different metabolic challenges remains to be determined. *In vitro* glucose-stimulated insulin secretion from islets isolated from single Aab+ donors is preserved (72), but a case report of a GADA+ donor showed fasting blood glucose and HbA1C in prediabetic ranges and marked glucose intolerance (50). Studies have also shown that individuals at greater risk of developing T1D fail to increase insulin/C-peptide secretion with age and have diminished first-phase insulin response to intravenous glucose (73). Whether islet vascular dysfunction potentiates this functional decline, which accelerates as the disease approaches (74), will be the focus of future work.

### Limitations of the study

In this study, we have not identified the molecular mechanisms underlying hemodynamic changes in islets. One of the reason is that we still know very little how islet cells (endocrine, vascular, immune cells etc) work together to maintain homeostasis and proper function. Understanding how different cells in the human islet communicate with each other under normal physiological conditions is crucial to be able to later determine what becomes compromised in a disease state. Another limitation of the study is that the limited amount of living human material we received prevented us from correlating our functional findings with the presence of residual (insulin-positive) beta cells. However, recent studies have shown that a considerable amount of islets still contain beta cells (∼40-60%) at diagnosis in people who developed T1D as teenagers or later (75), which is the case of the donors used in this study. We are also aware that only a subset of single Aab+ donors represent true prediabetic individuals who would have developed T1D. Unfortunately, we have not received living pancreas slices from donors with multiple Aab+ to functionally examine the islet microvasculature at later stages of islet autoimmunity.

## Acknowledgements

The authors would like to thank Drs. Alberto Pugliese (City of Hope, CA), René Barro-Soria, Ruy Andrade Louzada (University of Miami, Fl) and Profs. Nilda Gallardo and Antonio Andres Hueva (Castilla La Mancha University) for careful revision and discussion of the manuscript. The authors would like to thank the Network for Pancreatic Organ Donors with Diabetes (nPOD), in particular the organ donors, their families and the nPOD slicing team, under Dr. Irina Kusmartseva’s supervision, for producing living pancreas slices that allowed us to conduct these studies. This manuscript used data acquired from the Human Pancreas Analysis Program (HPAP-RRID:SCR_016202) Database (https://hpap.pmacs.upenn.edu), a Human Islet Research Network (RRID:SCR_014393) consortium (UC4-DK-112217, U01-DK-123594, UC4-DK-112232, and U01-DK-123716).

## Funding

This work has been funded by NIH grants K01DK111757 and R01DK133483 (to J.A.), by NIDDK-supported Human Islet Research Network (HIRN, RRID:SCR_014393; https://hirnetwork.org; UC4 DK104162) New Investigator Pilot Award (to J.A.) and by the Helmsley foundation Pilot Award for nPOD team science (AWD-007061; to J.A.).

## Author contributions

L.M.G performed immunohistochemistry, functional imaging with living human pancreas slices, and quantified data; M.M.F.Q performed bioinformatic analysis of scRNAseq data; M.B. analyzed immunohistochemical data; M.M. analyzed functional data using MatLab; J.A. designed the study, analyzed data and wrote the manuscript. All authors discussed the data and revised the manuscript. Further information and requests for resources or reagents should be directed to the Lead Contact, Joana Almaça.

## STAR material and methods

### Living human pancreas slices

We obtained human living pancreas slices (150 μm thickness; pancreas pieces were taken from the tail of pancreas) from de-identified cadaveric donors from the Network of Pancreatic Organ Donors with Diabetes (nPOD). Slices were produced by nPOD and 9-11 slices/donor were shipped overnight from Gainesville to Miami and used within 4-36h after arrival. Slices were cultured as previously described (76). Briefly, upon arrival, living slices were placed on perfluorocarbon (PFC) membrane AirHive cell culture dishes containing BrainPhys neuronal medium (Stemcell Technologies, cat. nr. 05790) supplemented with 2% B27 minus-insulin (Invitrogen, cat. nr. A1895601), 1% penicillin-Streptomycin-Amphotericin B solution (Sigma Aldrich, cat. nr. A5955), 1% Glutamax (Invitrogen, cat. nr. 35050061), 5.5 mM D-glucose (Sigma Aldrich, cat. nr. G8644), 100 µg/ml trypsin inhibitor from Glycine max (Sigma Aldrich, cat. nr. T6522), 10 µg/ml aprotinin (Sigma Aldrich, cat. nr. A6106), 10 µg/ml chymostatin (solubilized initially in DMSO; Sigma Aldrich, cat. nr. 11004638001) and 1% HEPES buffer (Invitrogen, cat. nr. 15630080). Dishes containing 3-4 slices and culture medium were placed in a humidified incubator at 30 °C, and meedium was changed every 12 h.

In this study, we received living pancreas slices that we used for physiology and/or immunohistochemistry from 12 Aab-, non-diabetic donors (ND), 7 single autoantibody positive (GADA+) donors and 9 T1D organ donors, with T1D duration of 0-4 years. Donor age ranged from 12-33 years old, and from both genders (details on donor characteristics are included in Table S1). All T1D donors were positive for islet autoantibodies (Table S1). C-peptide levels and glycated hemoglobin values of different organ donors used in this study are shown in Figure S1.

### Confocal imaging of living pancreas slices

Living human pancreas slices were incubated with the cytosolic calcium indicator ([Ca^2+^]i) Fluo4-AM (6 μM, Invitrogen, cat. nr. F14201) in 3 mM glucose solution prepared in HEPES buffer (125 mM NaCl, 5.9 mM KCl, 2.56 mM CaCl_2_, 1 mM MgCl2, 25 mM HEPES, 0.1% BSA [w/v], pH 7.4), supplemented with aprotinin (25 KIU, MilliporeSigma, cat. nr. A6106), at room temperature and in the dark with either (1) DyLight 649 lectin from *Lycopersicon Esculentum* (3.3 mg/mL, VectorLabs, cat. nr. DL1178) for 1 hour; or with (2) a fluorescent-conjugated antibody against the pericyte marker neuron-glial antigen 2 (NG2-alexa647; 1:50, R&D Systems, cat. nr. Fab2585R) for 2 hours. After incubation, living pancreas slices were placed on a coverslip in an imaging chamber (Warner instruments, Hamden, CT, USA) and imaged under an upright confocal microscope (Leica TCS SP8 upright; Leica Microsystems). The chamber was continuously perfused with HEPES-buffered solution containing 3 mM glucose and confocal images were acquired with LAS AF software using a 40X water immersion objective (NA 0.8). We used a resonance scanner for fast image acquisition to produce time-lapse recordings spanning 50-100 μm of the slice (z-step: 5-10 μm, stack of 10-15 confocal images with a size of 512×512 pixels) at 5 seconds resolution (xyzt imaging). Fluo-4 fluorescence was excited at 488 nm and emission detected at 510–550 nm, DyLight 649 lectin or NG2-alexa647 antibody were excited at 638 nm and emission detected at 648-690 nm.

We recorded changes in islet pericyte [Ca^2+^]i and capillary diameter induced by exchanging extracellular glucose concentration from 3 mM to 11mM (3G to 11G; 11G applied for 5-7 min), norepinephrine (20 μM; applied for 4 min in 3G), endothelin-1 (10 nM; applied for 5 min in 3G), angiotensin II (100 nM; applied for 4 min in 3G), adenosine (50 μM; applied for 4 min in 3G), bosentan (100 nM; applied for 5 min in 3G) and theophylline (20 μM; applied for 5 min in 11G). Endocrine cell [Ca^2+^]i responses to high glucose (11G) and KCl depolarization (25 mM; for 2 min in 3G) were also recorded. Islets in slices were identified using the backscatter signal produced by dense-core granules. To quantify changes in pericyte [Ca^2+^]i, we drew regions of interest around NG2-alexa647 labeled cells in islets as previously described (23). We quantified changes in mean Fluo4 fluorescence intensity using ImageJ software (http://imagej.nih.gov/ij/). Changes in fluorescence intensity were expressed as percentage over baseline (ΔF/F). The baseline was defined as the mean of the first 10 values of the control period of each recording [basal glucose concentration (3 mM) for all stimuli, except for theophylline for which baseline values were calculated during 11 mM glucose]. After subtracting the baseline, we calculated the net area under the curve of fluorescence traces to estimate the magnitude of the effect of each stimulus on cellular [Ca^2+^]i. MatLab software (https://www.mathworks.com/) was used to generate heatmaps showing changes in fluorescence of all recorded pericytes.

### Vasomotion analysis

Blood vessels in living slices were labeled with DyLight-649 fluorescent lectin. We quantified changes in vessel diameter as previously described (23). Briefly, we drew a straight-line transversal to the blood vessel borders and used the “reslice” z-function in ImageJ to generate a single image showing the changes in vessel diameter over time (xt scan; temporal projection; Figure 3A). xt scan (resliced) images were despeckled, blurred with Gaussian filter sigma = 1, before enhancing contrast, and image sharpened using the following kernel in order to emphasize vertical lines: −10 −5 50 −5 −10; −10 −5 75 −5 −10; −10 −5 50 −5 −10. An horizontal line was drawn on the xt scan (resliced) image (which corresponds to a single time point), and an array of pixel intensity values was sorted and first 2 maxima were considered vessel borders (tolerance = 4). Vessel diameter was calculated by subtracting these 2 values. For each stimulus, an average diameter value was calculated using the 10 last diameter values obtained during stimulus application. To determine the extent of constriction/dilation, we pooled diameter data from different capillaries from different islets for each group of donors and calculated the relative change in diameter (as % of baseline (3G) vessel diameter).

### Immunohistochemistry

After physiological recordings, living human pancreas slices from non-diabetic, Aab+ and T1D organ donors were fixed for 1h with 4% PFA, washed 3x in PBS, and stored at 4’C. For immunohistochemistry, slices were incubated in blocking solution (PBS-Triton X-100 0.3% and Universal Blocker Reagent; Biogenex, San Ramon, CA) for 3h. Thereafter, slices were incubated for 48h (20°C) with primary antibodies diluted in blocking solution. To identify pancreatic islets in slices, we immunostained either beta cells (insulin, 1:5) or delta cells (somatostatin; 1:250). We labeled pericytes using an anti-NG2 antibody (1:50-1:100), smooth muscle cells/myofibroblasts using anti-*α*SMA (1:250), endothelial cells using anti-CD31 (1:25), sympathetic nerve fibers using anti-tyrosine hydroxylase (TH; 1:400), basement membrane using anti-collagen IV antibody (1:250), periostin (1:250). Information on the different antibodies used is provided as Supplementary material (Table S2). Immunostaining was visualized using conjugated secondary antibodies (1:500 in PBS; 16h at 20°C; Invitrogen, Carlsbad, CA). Cell nuclei were stained with dapi. Slides were mounted with Vectashield mounting medium (Vector Laboratories) and imaged on an inverted laser-scanning confocal microscope (Leica TCS SP5; Leica Microsystems) with LAS AF software using a 63X oil immersion objective (NA 1.4). Image analysis was performed using artificial intelligence and automated macros. Briefly, to quantify the density of each protein/marker in islets, an hormone staining (insulin or somatostatin) was used to locate islets within the image, while DAPI staining was used to outline the whole tissue area. The image was then segmented into islet area and exocrine tissue area, and the immunostained marker was thresholded based on mean image value for that channel. A mask for each immunostaining was created and the corresponding area was measured. To estimate islet *versus* acinar densities, we divided the immunostained area for each marker by the corresponding tissue area. To quantify colocalization between collagen IV and the pericyte marker NG2, we calculated Mander’s coefficients in confocal images of islets from different donors using the ImageJ plugin “JACoP: Just Another Co-localization Plugin”.

### Single-cell RNAseq analysis

To examine changes in gene expression related to the vasculature of different cell populations in the human pancreas, we used accessed the single cell RNAseq dataset (PancDB) from the Human Pancreas Analysis Program (HPAP; (37); https://hpap.pmacs.upenn.edu). This dataset contains 222,077 cells across 67 individuals. We chose human donors that matched the characteristics of those we had used for physiological experimentation (Table S1 and Figure S1). Data were subsetted from 20 individuals accounting for 7 single Aab+ (GADA+; 16,784 cells), 9 non-diabetic (17,603 cells) and 4 type-1 diabetic donors (disease duration: 0-3 years; 8,813 cells), collectively represented by 43,200 cells. We employed a comprehensive analytical pipeline using custom designed R scripts similar to that outlined previously (77). Cells clusters were selected based on their specific expression of different gene sets. Briefly, beta cells were identified based on the expression of *INS, IAPP, PDX1* and *MAFA*, alpha cells on *GCG, DPP4* and *GC* genes, endothelial cell on *VWF* and *ENG* genes and stellate cells (which include pericytes) on the expression of *PDGFRB* and *COL1A1* genes (Figure S5). The complete analytical architecture is deposited in Github (https://github.com/jalmaca/Microvasculature_T1D). In this analysis we consider a differentially regulated gene when change in gene expression (fold change (FC) values) were higher than 1.2 (upregulated gene; indicated with an *) or below 0.8 (downregulated gene; indicated with an *’), having a *p-value* less than 0.05 and being expressed in at least in 25% of cells across a cluster.

### Statistical Analyses

For statistical comparisons we used Prism 9 (GraphPad software, La Jolla, CA) and performed one-way ANOVA followed by Tukey’s multiple comparisons test when comparing data obtained from non-diabetic donors, single Aab+ and T1D organ donors. Paired t-tests were used to determine if changes in diameter induced by a certain stimulus for each individual capillary were significantly different from baseline values. One sample t tests were used to determine if average relative changes in vessel diameter induced by different stimuli were significantly different from 0. *p* values < 0.05 were considered statistically significant (indicated with an * in figures). Throughout the manuscript we present data as mean ± SEM.

## Notes

https://github.com/jalmaca/Microvasculature_T1D

### Competing Interest Statement

The authors have declared no competing interest.

