## Supplementary information for "Pericyte dysfunction and impaired vasomotion are hallmarks of islets during the pathogenesis of type 1 diabetes"

### Supplementary Tables

**Supplementary Table 1.** Characteristic of organ donors used in the study. Living pancreas slices were obtained from nPOD (Gainesville, Florida) and used for physiology and/or for immunohistochemistry.

| Case ID | Donor Type | Gender | Age | Race | BMI | AutoAb | T1D (yrs) | HbA1c | C-peptide | Physiology | IHC | scRNAseq |
|---|---|---|---|---|---|---|---|---|---|---|---|---|
| 6516 | ND | M | 20 | Caucasian | 28.8 | | | 5.5 | 8.91 | x | | |
| 6531 | ND | F | 19 | Hispanic | 30 | | | 5.5 | 26.53 | x | x | |
| 6535 | ND | F | 31 | Caucasian | 29.6 | | | | 6.59 | x | | |
| 6537 | ND | M | 33 | Caucasian | 20.8 | | | 5.6 | 0.32 | x | x | |
| 6539 | ND | M | 24 | Hispanic | 19.3 | | | 5.7 | 39.23 | x | x | |
| 6546 | ND | M | 22 | Asian | 23.7 | | | 5.6 | 11 | x | x | |
| 6548 | ND | M | 20 | Caucasian | 23.8 | | | 5.7 | 4.04 | x | x | |
| 6552 | ND | F | 33 | Caucasian | 21.9 | | | 5.6 | 1.8 | x | | |
| 6555 | ND | F | 18 | Caucasian | 18.4 | | | 5.2 | 3.6 | x | x | |
| 6468 | ND | M | 16 | Caucasian | 15.9 | | | 4.4 | 5.27 | | x | |
| 6470 | ND | M | 30 | Caucasian | 22.5 | | | 5.5 | 6.03 | | x | |
| 6471 | ND | M | 16 | African American | 18.4 | | | 5.5 | 1.99 | | x | |
| HPAP026 | ND | M | 24 | Caucasian | 20.8 | | | 4.9 | 0.25 | | | x |
| HPAP034 | ND | M | 13 | Caucasian | 18.6 | | | 5.2 | 12.7 | | | x |
| HPAP035 | ND | M | 35 | Caucasian | 26.9 | | | 5.2 | 15.9 | | | x |
| HPAP036 | ND | F | 23 | Caucasian | 16 | | | 5.2 | 1.12 | | | x |
| HPAP037 | ND | F | 35 | Caucasian | 21.9 | | | 5.3 | 4.75 | | | x |
| HPAP040 | ND | M | 35 | Caucasian | 23.9 | | | 5.4 | 7.01 | | | x |
| HPAP047 | ND | M | 8 | Caucasian | 16.8 | | | N/A | 1.24 | | | x |
| HPAP082 | ND | M | 25 | Caucasian | 23.9 | | | 5.6 | 2.7 | | | x |
| HPAP099 | ND | F | 28 | Hispanic | 24.7 | | | 5 | 6.62 | | | x |
| 6532 | AAB+ | M | 20 | Hispanic | 23.9 | GADA+ | | 5.9 | 22.12 | x | x | |
| 6538 | AAB+ | M | 19 | Caucasian | 32.9 | GADA+ | | 5.8 | 11.33 | x | x | |
| 6558 | AAB+ | F | 21 | African American | 27.8 | GADA+ | | 4.4 | 8.03 | x | x | |
| 6562 | AAB+ | F | 29 | Caucasian | 18.2 | GADA+ | | 5.6 | 13.19 | x | x | |
| 6569 | AAB+ | F | 20 | Hispanic | 24.2 | GADA+ | | 6 | 2.44 | x | x | |
| 6573 | AAB+ | F | 24 | Caucasian | 35.5 | GADA+ | | 5.1 | 1.85 | x | x | |
| 6575 | AAB+ | M | 23 | Caucasian | 26.9 | GADA+ | | 5.5 | 5.21 | x | x | |
| HPAP024 | AAB+ | M | 18 | Caucasian | 24.4 | GADA+ | | 5.5 | 5.6 | | | x |
| HPAP029 | AAB+ | M | 23 | Caucasian | 28.6 | GADA+ | | 5.3 | 3.83 | | | x |
| HPAP038 | AAB+ | M | 13 | Caucasian | 18.3 | GADA+ | | 5.7 | 8.29 | | | x |
| HPAP045 | AAB+ | F | 27 | Caucasian | 26.2 | GADA+ | | 5.2 | 1.7 | | | x |
| HPAP050 | AAB+ | F | 21 | Hispanic | 28.9 | GADA+ | | 5.1 | 3.79 | | | x |
| HPAP072 | AAB+ | M | 19 | Hispanic | 23.1 | GADA+ | | 5.6 | 4.37 | | | x |
| HPAP092 | AAB+ | M | 21 | Hispanic | 25.6 | GADA+ | | 5.6 | 15.35 | | | x |
| 6523 | T1D | F | 12 | African American | 22.5 | GADA+ mIAA*+ | 3 | 11.1 | 0.04 | x | | |
| 6536 | T1D | F | 20 | Caucasian | 25.4 | GADA+ | 4 | 12.7 | 0.04 | x | x | |
| 6550 | T1D | M | 25 | Caucasian | 16.4 | GADA+ ZnT8A+ | 0 | 14 | <0.02 | x | x | |
| 6551 | T1D | M | 20 | Caucasian | 23.1 | GADA+IA2A+ mIAA+* ZnT8A+ | 0.58 | 6.4 | 0.11 | x | x | |
| 6563 | T1D | F | 14 | Caucasian | 25.5 | IA2A+ | 0 | 9.6 | 1.04 | x | x | |
| 6566 | T1D | M | 15 | Caucasian | 21.8 | GADA+IA2A+ mIAA+* ZnT8A+ | 2 | 10.3 | 0.05 | x | x | |
| 6456 | T1D | F | 30 | African American | 30.1 | GADA+ ZnT8A+ | 0 | 6.8 | 10.33 | | x | |
| 6469 | T1D | F | 27 | Caucasian | 26.9 | GADA+ | 1.5 | 7.4 | 0.66 | | x | |
| 6480 | T1D | M | 17 | Caucasian | 27.1 | IA2A+ mIAA*+ | 2.5 | 10.2 | 0.13 | | x | |
| HPAP032 | T1D | F | 10 | Caucasian | 16.3 | IA2A+ mIAA+ | 3 | 9 | 0.02 | | | x |
| HPAP064 | T1D | M | 24 | African American | 16.9 | ZnT8A+ | N/A | 13 | 0.25 | | | x |
| HPAP071 | T1D | F | 12 | African American | 15.4 | IA2A+ mIAA+ | 3 | 9.8 | 0.06 | | | x |
| HPAP084 | T1D | F | 12 | African American | 18.5 | GADA+ IA2A+ mIAA+ | N/A | 13.3 | 2.2 | | | x |

**Supplementary Table 2.** Antibodies and reagents used in the study.

| **Antibodies** | **Company** | **Catalog number** |
|---|---|---|
| Neuron-glial antigen 2 (NG2) | R&D | FAB2585R |
| Alpha smooth muscle actin (aSMA) | Sigma | A5228 |
| PECAM (CD31) | Millipore | 550389 |
| Tyrosine hydroxylase (TH) | Millipore | AB152 |
| Collagen IV | Abcam | AB6586 |
| Periostin | R&D | AF2955 |
| Insulin | Dako | IR002 |
| Somatostatin | Millipore | MAB354 |
| **Chemicals and peptides** | | |
| Theophylline | Sigma | T1633 |
| Norepinephrine | Tocris | 5169 |
| Endothelin 1 | Sigma | E7764 |
| Angiotensin II | Tocris | 1158 |
| Bosentan | Sigma | 1265 |
| *Lycopersicon esculentum* Lectin | Vector Labs | DL-1178-1 |

**Supplementary Figures**

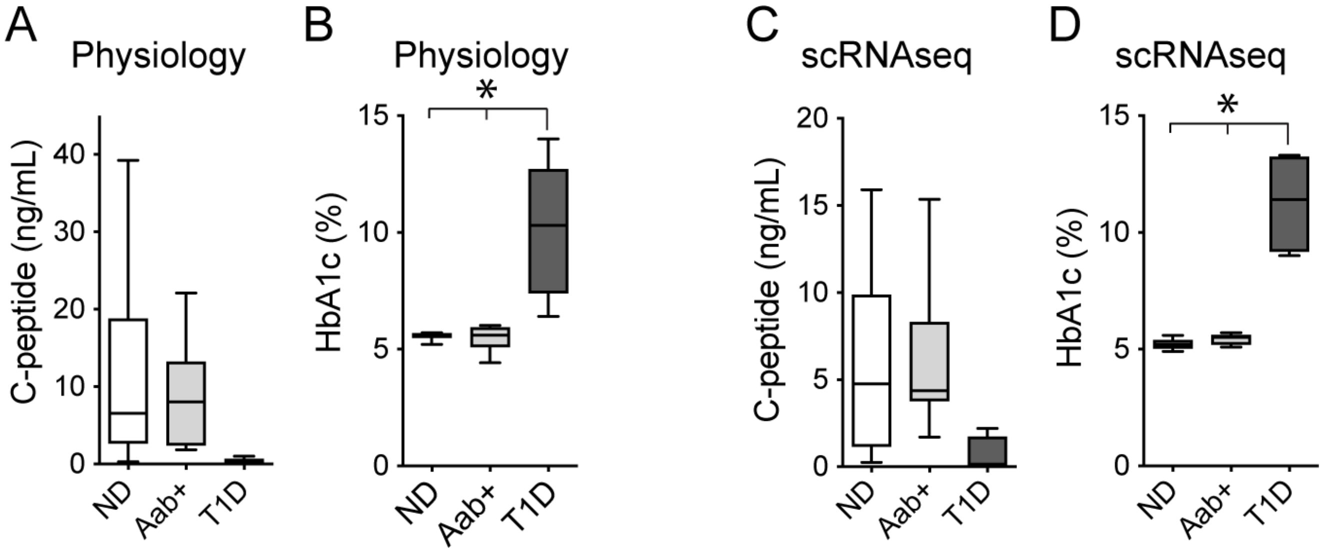

**Supplementary Figure 1. C-peptide and HbA1C levels of organ donors used in this study.** (A,B) Plasma C-peptide (A) and HbA1C levels (B) of organ donors whose pancreas slices were used physiology in this study (non-diabetic donors (ND), GADA+ donors (Aab+) and T1D donors). * indicates $p < 0.05$ when compared with levels of ND and Aab+ donors (one-way ANOVA followed by Tukey's multiple comparisons test). (C,D) Plasma C-peptide (C) and HbA1C levels (D) of organ donors chosen from HPAP database to analyze changes in gene expression in pancreatic endocrine alpha and beta cells, endothelial and stellate cells.

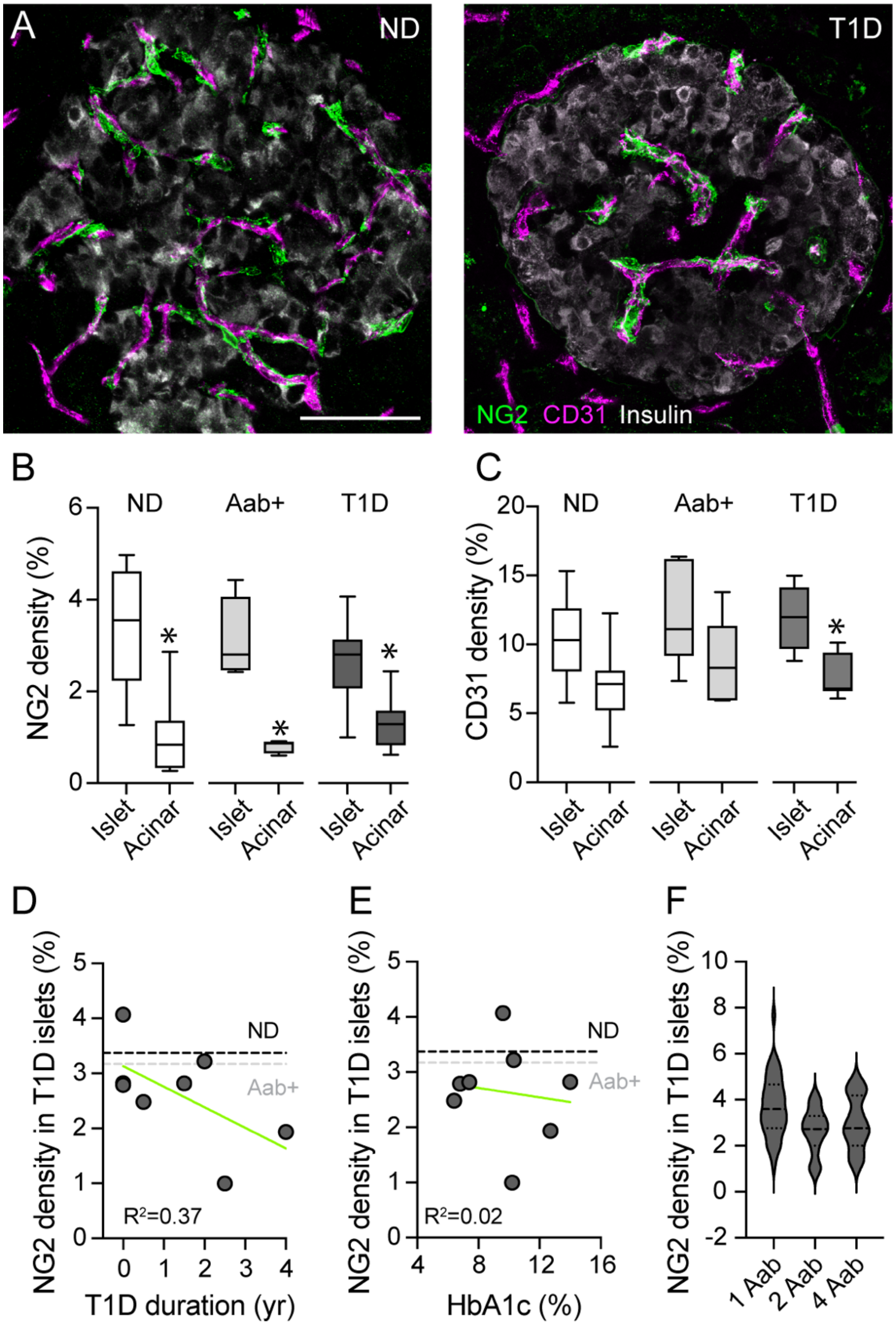

**Supplementary Figure 2. Changes in pericyte and endothelial cell densities in islets from Aab+ and T1D organ donors.** (A) Maximal Projections of confocal images of islets in fixed pancreatic slices from a non-diabetic (ND; nPOD6531) and a T1D donor (T1D duration 1.5y; nPOD6469) immunostained with antibodies against the endothelial cell marker CD31 (magenta), the pericyte marker neuron-glial antigen 2 (NG2; green) and insulin (gray). Note that in T1D islets several capillaries lack pericyte coverage. Scale bar = 50 μm. (B,C) Quantification of the % of tissue area (islet or acinar) immunostained with NG2 (B) or CD31 (C)

in pancreas slices from non-diabetic (ND, n=9 donors), GADA+ (Aab+, n=7 donors) and T1D donors (T1D, n=8). For each donor, around 5-7 islets were imaged, and an average value was calculated. * indicates $p < 0.05$ when compared with islet densities (one-way ANOVA followed by Tukey's multiple comparisons test. (D,E) Correlation between the density of pericytes in T1D islets and the duration of T1D (D) or the HbA1C levels (E). For each donor, an average of 5-7 islets is shown. Linear regressions are not significant. Dashed lines indicate average NG2 densities obtained for non-diabetic (ND; black) and Aab+ donors (gray). (F) Density of pericytes in islets from T1D donors depending on the number of circulating autoantibodies. Data represent individual islets (n=37 (1 Aab+), 12 (2 Aab+) and 20 (4 Aab+)).

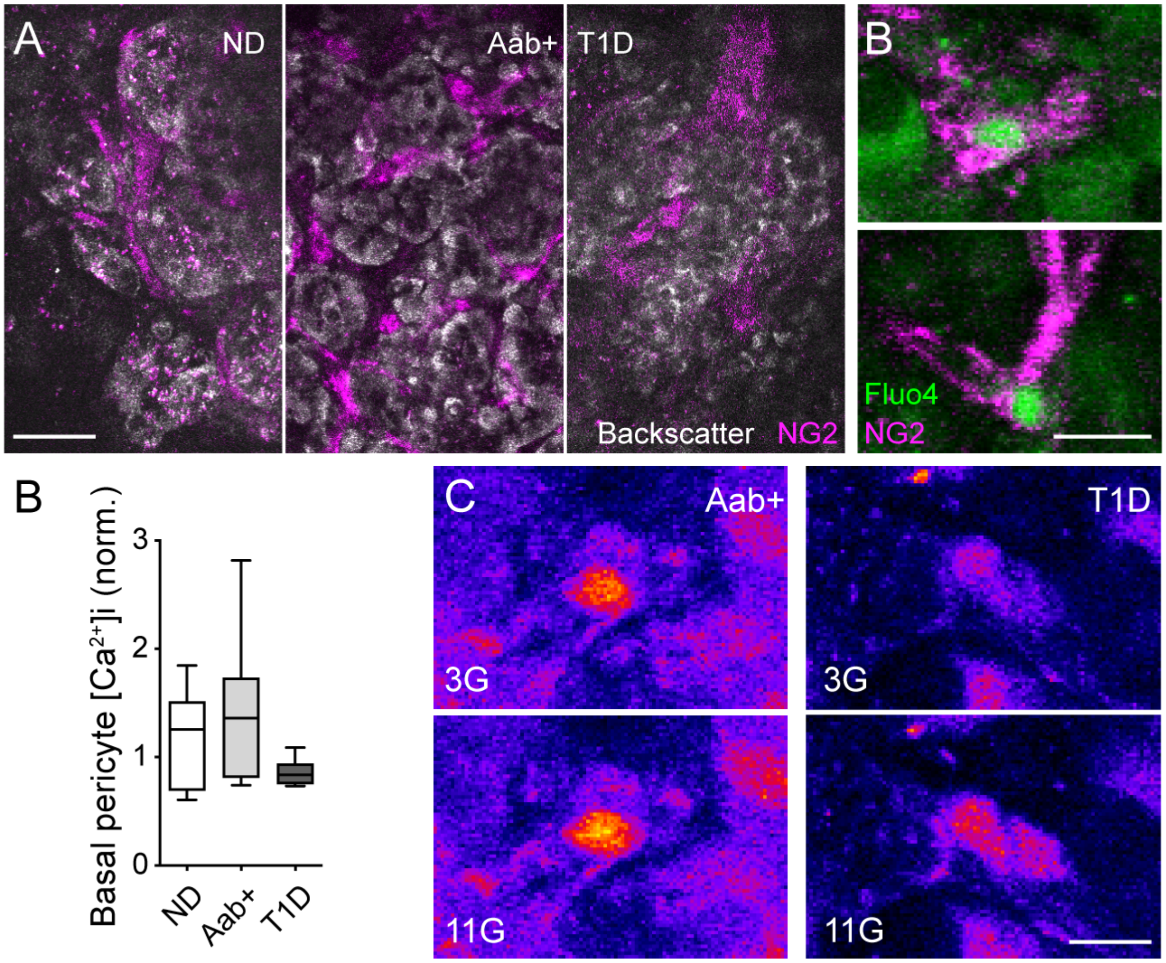

**Supplementary Figure 3. Using living pancreas slices to measure pericyte $[Ca^{2+}]i$ responses to high glucose.** (A) Confocal images of islets (backscatter; gray) in living pancreas slices from a non-diabetic (ND; nPOD6546), a Aab+ (nPOD6538) and a T1D donor (T1D duration 2y; nPOD6566) showing NG2-alexa647-labeled pericytes (magenta). (B) Slices were loaded with the calcium indicator Fluo4 (green). Different cells incorporate the indicator but pericytes can be distinguished as they are labeled with NG2-alexa647. (C) Quantification of basal $[Ca^{2+}]_i$ levels in islet pericytes normalized to fluorescence levels in the whole islet (n = 9 ND donors; 6 Aab+ donors; 6 T1D donors). 8-12 pericytes were analyzed per islet, 3-5 islets per donor and an average per donor was calculated. (C) Confocal images of pericytes in islets from Aab+ or T1D donors in low glucose (3G) or upon high glucose stimulation (11G). These pericytes are not inhibited by high glucose application. Scale bars = 20 μm (A), 10 μm (B,C).

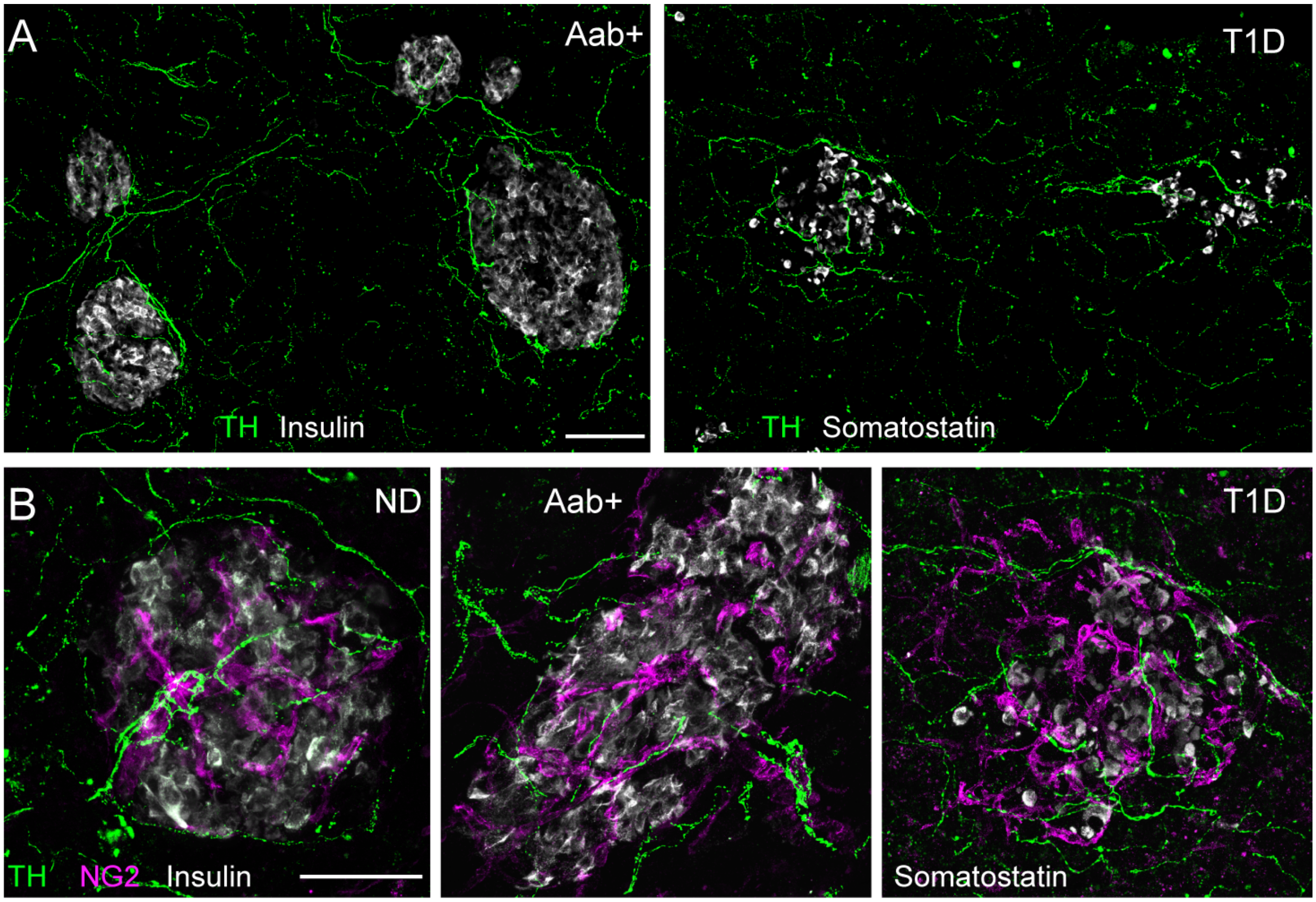

**Supplementary Figure 4. Sympathetic innervation patterns of islets at different stages of T1D.** (A) Maximal projections of confocal images of fixed pancreas slices from an Aab+ donor (nPOD6573) and a T1D donor (T1D duration 2 years; nPOD6566) immunostained with the sympathetic nerve marker tyrosine hydroxylase (TH; green). Islets were identified with insulin or somatostatin immunostainings (gray). Sympathetic nerves are present in both endocrine and exocrine compartments of the pancreas. (B) Projections of confocal images of fixed pancreas slices from a ND (nPOD6546), an Aab+ donor (nPOD6573) and a T1D donor (nPOD6566) immunostained for TH (green), insulin/somatostatin (gray) and the pericyte marker NG2 (magenta). Sympathetic nerves reach the islet and contact a subset of islet pericytes (~20% of NG2-expressing pericytes) within the islet parenchyma. Scale bars = 50 μm (A,B).

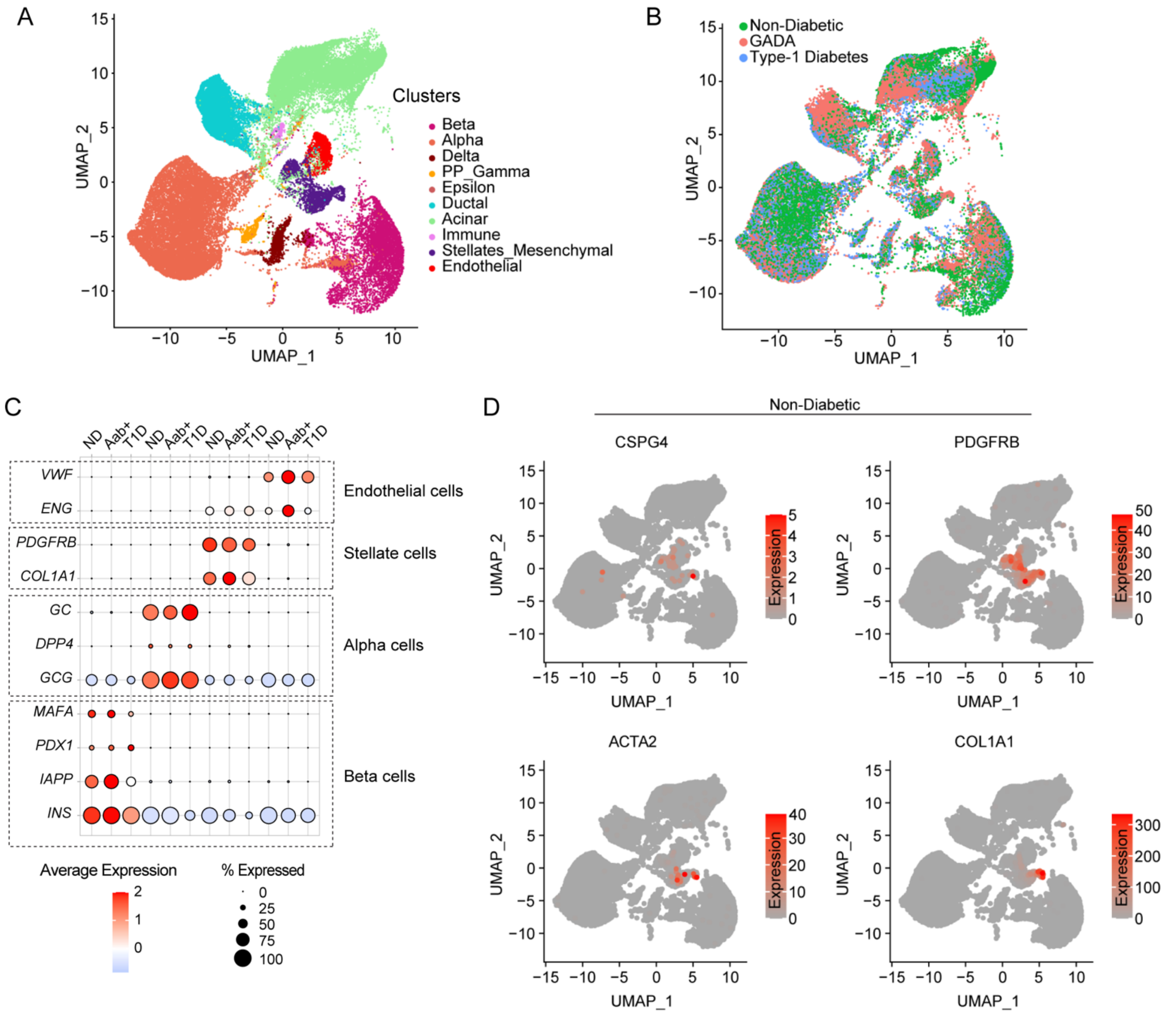

**Supplementary Figure 5. Identification of different cell populations in the human pancreas by scRNAseq.** (A) Uniform Manifold Approximation Projection (UMAP) of human pancreatic cells taken from the HPAP database and clustered based on cell type. Each dot represents the unique transcriptional profile of a single cell. (B) UMAP plot showing the cluster distribution of cells across three disease states: non-diabetic, single Aab+ (GADA+) and T1D. (C) Dotplot showing the expression of key genes specific to beta, alpha, stellate, and endothelial cell populations. Size of the circle denotes percentage of cells within a cluster expressing a gene and the intensity of the color denotes the magnitude of normalized average expression. (D) UMAP plots of gene expression in non-diabetic donors for genes *CSPG4, PDGFRB, ACTA2* and *COL1A1*.

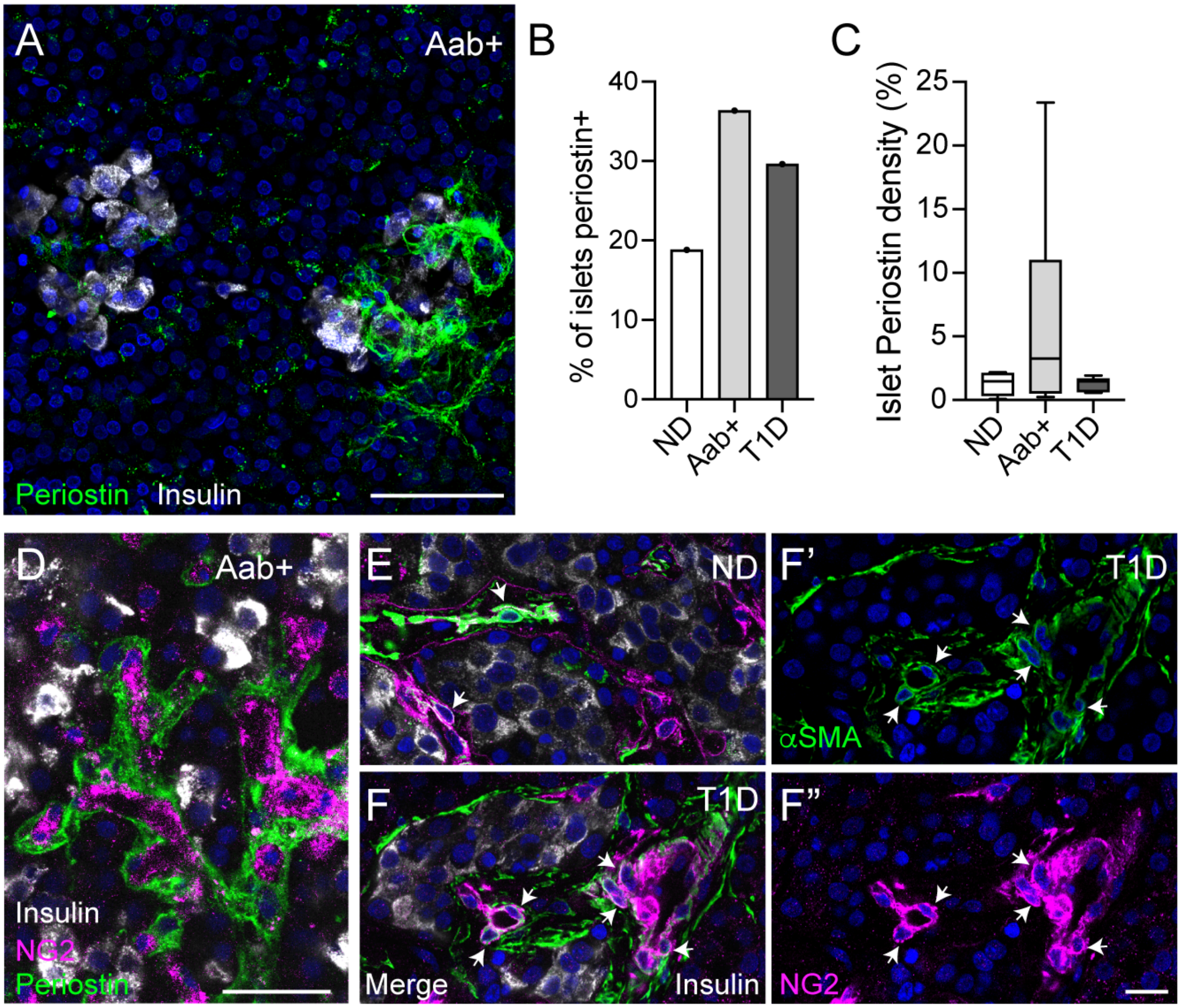

**Supplementary Figure 6. Pericytic expression of myofibroblast markers in islets at different stages of T1D.** (A) Maximal projection of confocal images of islets in a fixed pancreatic slice from an Aab+ donor (nPOD6538) immunostained for insulin (gray) and the ECM and myofibroblast marker periostin (green). Periostin accumulates in spaces between beta cells but its presence in islets is heterogenous. (B) Quantification of the proportion of islets for each group of donors that contain periostin. (C) Quantification of the % of islet area immunostained with periostin in pancreas slices from non-diabetic, Aab+ and T1D donors (n = 6-16 islets/group). (D) Projection of confocal images of an islet in a fixed pancreas slice from an Aab+ donor (nPOD6538) immunostained for insulin (gray), periostin (green) and NG2 (magenta). (E,F-F") Confocal images of regions within islets in sections from a ND donor (E) and a T1D donor (T1D duration 1.5 years; nPOD6469; F-F") showing pericytes (labeled with anti-NG2 antibody; in magenta) and αSMA (green). Beta cells are shown in gray (insulin). Note the increase in the proportion of islet pericytes that expresses both NG2 and αSMA (white arrows). Scale bars = 50 μm (A), 20 μm (D) and 10 μm (E,F-F").
